# Convergent genomic trajectories shape adaptation to life on land across animal lineages

**DOI:** 10.1101/2025.10.02.680026

**Authors:** Gemma I. Martínez-Redondo, Klara Eleftheriadi, Judit Salces-Ortiz, Nuria Escudero, Fernando Ángel Fernández-Álvarez, Belén Carbonetto, Carlos Vargas-Chávez, Raquel García-Vernet, Javier Palma-Guerrero, Libe Rentería, Iñaki Rojo, Cristina Chiva, Eduard Sabidó, Aureliano Bombarely, Rosa Fernández

**Affiliations:** Metazoa Phylogenomics and Genome Evolution Lab, Institute of Evolutionary Biology (CSIC-UPF), Barcelona (Spain); Universitat de Barcelona, Barcelona (Spain); Centro Oceanográfico de Gijón, Instituto Español de Oceanografía, Gijón (Spain); Institut de Ciències del Mar (ICM-CSIC), Barcelona (Spain); Research Institute of Organic Agriculture (FiBL), Aargau (Switzerland); Citizen scientist, Sopelana (Spain); Centre for Genomic Regulation (CRG), Barcelona Institute of Science and Technology (BIST), Barcelona (Spain); Universitat Pompeu Fabra (UPF), Barcelona (Spain); Instituto de Biología Molecular y Celular de Plantas, Valencia (Spain)

## Abstract

How animals repeatedly adapted to life on land is a central question in evolutionary biology. While terrestrialisation occurred independently across animal phyla, it remains unclear whether shared genomic mechanisms underlie these transitions. Here, we combine large-scale comparative genomics, machine learning, and multi-omics data, including proteomics and transcriptomics from stress experiments relevant to terrestrial environmental challenges in 17 species, to investigate the genomic basis of animal terrestrial adaptation. Gene co-expression networks reveal that genes relevant to stress are largely lineage-specific, yet converge in function through the co-option of gene families pre-dating terrestrialisation events. Phylogenomic and machine learning analyses support a dominant role for early-evolving genes, enriched in stress-related functions, paired with a higher gene loss than gain at terrestrialisation nodes. Our findings support a model of lineage-specific genomic changes involving mostly conserved genes that converged at the functional level during the independent transitions to terrestrial life.

## Introduction

The transition of animals to land (terrestrialisation) is one of the most transformative events in evolutionary history. Coming from relatively stable marine environments, animals underwent radical physiological and morphological adaptations to cope with the environmental challenges of life on land, including desiccation, higher oxygen concentration, ultraviolet radiation, and changes in osmotic pressure and chemical exposure^1–5^. As such, terrestrialisation represents not just a change in habitat, but a major functional reconfiguration of animal biology. Fossil evidence suggests the first land exploration by marine arthropod relatives occurred around 470 million years ago^4–6^, likely driven by the need to mate or lay eggs out of predators’ reach. Over millions of years, terrestrial habitats became progressively inhabited as environments became more hospitable due to the biota’s own influence^1,7,8^. Unlike plants (*sensu stricto*), which terrestrialised only a few times, animals (Metazoa) have independently transitioned to land several times across at least nine out of the 31 animal phyla (Fig. 1a,b). Terrestrialisation in animals is highly heterogeneous, with many species remaining semi-terrestrial, requiring water at specific life stages, while only a few complete their entire life cycles on land^10,11^. These transitions occurred via direct marine-to-terrestrial routes or freshwater intermediaries utilised by certain groups like platyhelminthes, molluscs, crabs, and vertebrates^12–15^ (Fig. 1c). This rich tapestry of terrestrialisation, characterised by diverse dependencies on water, varied environmental routes to land, and staggered timelines, underscores the multifaceted nature of this evolutionary achievement.

**Figure 1.**
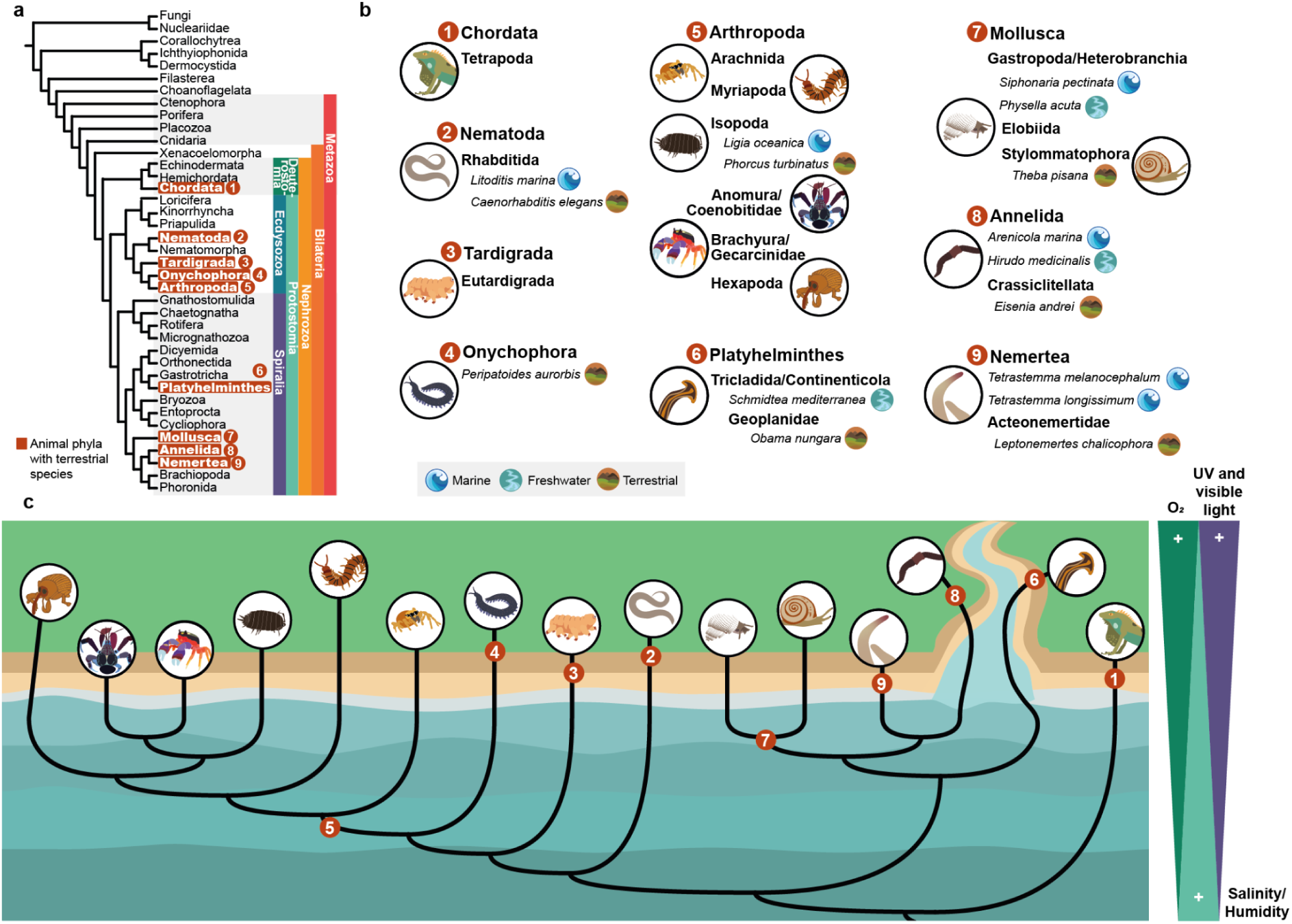
Animal terrestrialisation at a glance. **a**, Animal tree of life highlighting the phyla where there has been at least one terrestrialisation event. **b**, Main animal terrestrialisation events within animal phyla discussed in this work (in bold), with the species used in the stress experiments below. **c**, Simplified view of paths towards terrestrial life in animals, and the differences in some abiotic factors between habitats.

Most previous studies have focused on individual species^16,17^, specific phyla^18–21^ or particular gene families^22,23^, without assessing whether genomic adaptation to terrestrial life involved convergent genomic changes across all animal phyla, or arose through lineage-specific mechanisms. To address this, we conducted a broad comparative analysis of nearly 1,000 animal species, integrating genome-wide data with expression profiles from experiments that mimic environmental stressors faced during terrestrialisation^24–27,28,29^, along with proteomic data, in diverse aquatic and terrestrial invertebrates. We also identified gene families associated with habitat type through machine learning. By tracing the evolutionary origins and functions of these genes, we provide a comprehensive view of how animals have genomically adapted to life on land through repeated, but distinct, evolutionary routes.

## Results

### A kingdom-wide reconstruction of animal gene repertoire evolution

To understand the evolutionary foundations of animal terrestrialisation, we first reconstructed the origin and diversification of gene families across the animal kingdom. We performed orthology inference on high-quality protein-coding gene sets from 973 species (960 animals, Supplementary Data S1), spanning all 31 recognised animal phyla, resulting in over 500,000 orthogroups. Then, we dated the evolutionary origin of each orthogroup by identifying the most recent common ancestor of all the species in which that orthogroup is present and quantified the general patterns of gene gain (first appearance of a given orthogroup, as distinct from gene duplications) and subsequent loss across the metazoan tree (Fig. 2a, Supplementary Fig. S1). Our results showed that most orthogroups originated early in animal evolution, with many orthogroups established during the initial diversification of Metazoa, as previously proposed^30–32^. Notably, and contrary to previous research, we identified a striking burst of gene gain at the base of Protostomia (the lineage that includes all invertebrates that later transitioned to terrestrial environments), probably unveiled by our higher taxonomic sampling (Fig. 2b). In contrast, gene losses were much lower at deeper nodes, followed by independent bursts of losses, which are gradually reduced towards the tips of the tree. These patterns are also reflected in the gene gain-to-loss ratio (Supplementary Fig. S2), where there is a shift from higher gene gain to higher gene loss after Protostomia. While this bias may partly reflect methodological limitations, as gene gain/loss inference under parsimony is sensitive to tree topology, known major phylogenetic discordances in our tree generally predate terrestrialisation events and are unlikely to affect the overall pattern around these transitions. All in all, our results are congruent with previous findings^30–32^, and consistent with a biphasic model of evolution^33^.

**Figure 2.**
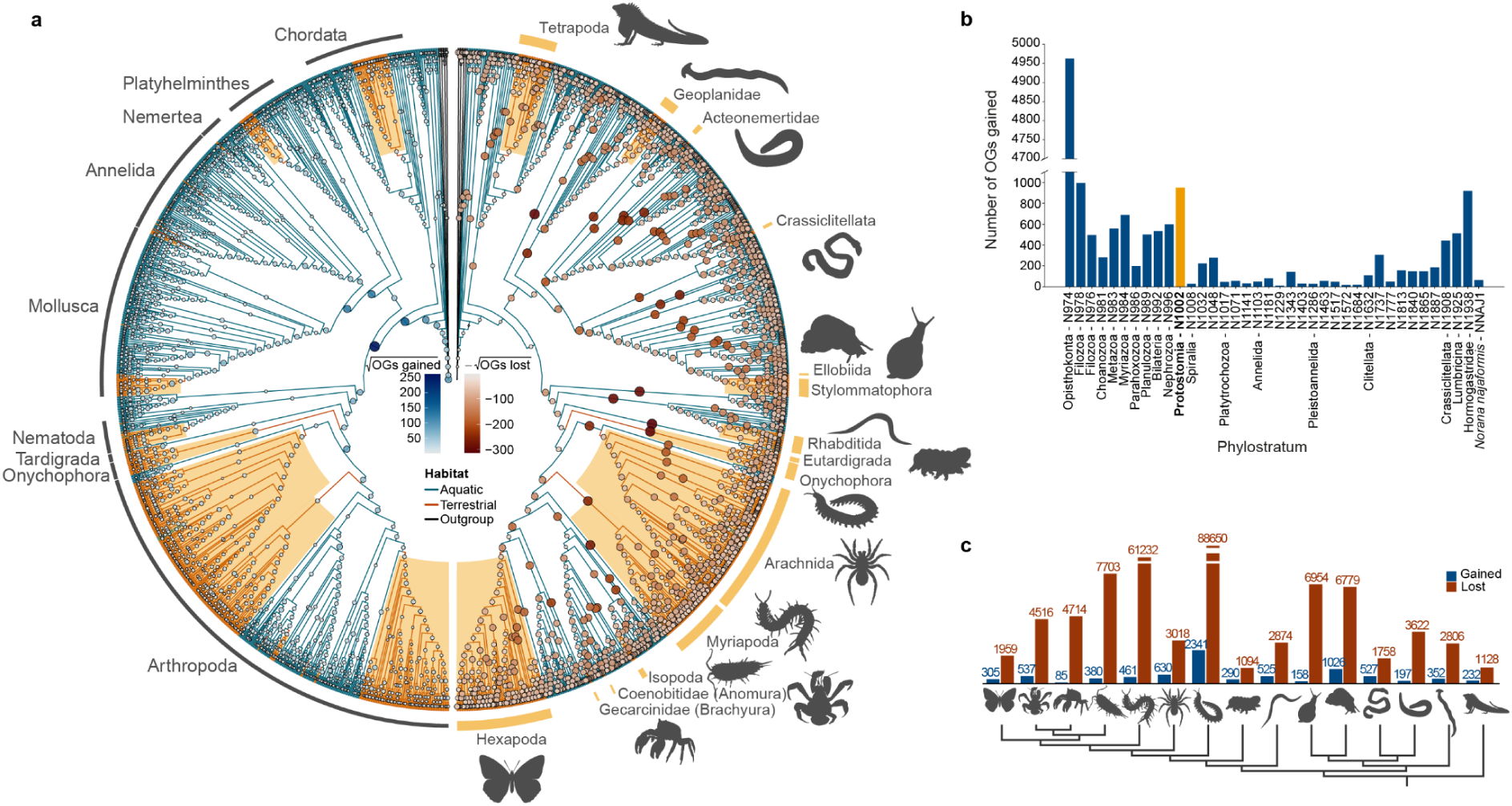
Orthogroup evolution across the animal kingdom. **a**, Mirror phylogeny of 960 animal species plus outgroups. Circles at each internal branch represent the square-root-scaled number of orthogroups gained (left, blue) and lost (right, brown). Terrestrial lineages are highlighted in yellow; the right phylogeny labels individual terrestrial lineages, while the left phylogeny indicates the phyla containing those lineages. Exact numbers of OGs gained and lost can be found in Supplementary Fig. S1. **b**, Phylostratigraphic origin of all genes in all nodes in a representative species (the earthworm *Norana najaformis,* Annelida), highlighting the gene gain burst in Protostomia (in yellow). **c**, Comparison of gene gain and loss across the 15 terrestrial clades with independent terrestrialisation events.

Our extensive dataset allowed us to study 15 independent animal terrestrialisation events represented by two or more species (Fig. 1b), encompassing both ancient transitions deep in the metazoan tree and more recent, lineage-specific events closer to the tips of the phylogeny (i.e., Tetrapoda, Geoplanidae, Acteonemertidae, Crassiclitellata, Ellobiida, Stylommatophora, Rhabditida, Eutardigrada, Onychophora, Arachnida, Myriapoda, Isopoda, Coenobitidae, Gecarcinidae, and Hexapoda). Compared to older nodes, fewer orthogroups originated at the nodes corresponding to terrestrial lineages. Instead, these nodes were marked by extensive gene loss, in some cases exceeding gene gains by more than an order of magnitude, with gene-to-loss ratios lower than 1 (Fig. 2c, Supplementary Fig. S2, Supplementary Data S2). This pattern was held consistently across independent terrestrialisation events in and within different phyla. To gain insight into the function of those genes, we then performed separate GO enrichment analysis on orthogroups inferred to be gained or lost at those nodes, using the full genomic gene set of representative species as background (Supplementary Figs. S3-92, see Methods). Genes belonging to both gained and lost orthogroups were significantly enriched in functional terms associated with response to stimuli (such as UV radiation or oxygen), DNA repair and maintenance, reproduction and meiosis, cell division and cell cycle regulation, reproduction, immunity and defense mechanisms, sensory perception and signaling, transport and homeostasis, and developmental processes related to organ formation (e.g., fin development GO term was enriched in lost orthogroups in tetrapods, Supplementary Fig. S90) and environmental adaptation. These functions are exemplified in the gene names that could be obtained using BLASTp. For example, some genes inferred to be gained in tetrapods include the human genes *HEY1*, *AMOT*, *LDHA, COL1A1, TNNT2,* and *TNNT3,* which are related to muscular and cardiovascular function and development^34–39^, and might have contributed to the locomotion changes during their water-to-land transition. Moreover, GO terms showed moderate functional convergence (measured as semantic similarity between the GO terms; see Methods) between independent terrestrial lineages (Supplementary Fig. S93a,b). For instance, when we compared hexapods and tetrapods, we found GO terms enriched in functions related to skin protection: cuticle development for hexapods (Supplementary Fig. S3) and keratinization for tetrapods (Supplementary Fig. S87). In addition, some OGs independently gained or lost in at least two terrestrial lineages contain genes with BLASTp hits to the same superfamilies, some of which having been previously linked to terrestrialisation^18–22,40^, including CYP450s, mucins, solute carrier proteins, aquaporins, vitellogenin, odorant receptors, or cuticle proteins, among others with more general cellular functions. This suggests a mechanism of paired gain and loss of genes with overlapping functions, reshaping the gene repertoire during terrestrialisation. Overall, the predominance of gene loss over gain at key phylogenetic nodes indicates that, during the transition to land, gene repertoire evolution may have relied more on the removal of functionally redundant or environmentally dispensable genes, facilitated by mutational robustness, rather than relying primarily on de novo gene innovation.

### Lack of a universal stress-response genomic toolkit for animal terrestrialisation

The transition from aquatic to terrestrial environments exposed animals to a consistent set of abiotic stressors, including water loss and osmotic imbalance, fluctuating oxygen availability, increased ultraviolet (UV) radiation and novel light regimes, as well as requiring changes in chemical signal detection, all of which are widely recognised as major selective pressures during terrestrialisation^24–29^. These stressors are expected to act across phylogenetically diverse lineages, making them suitable candidates for identifying shared or convergent genomic responses arising from the repeated and independent transition to terrestrial environments by animals.

To test whether a common stress-response genomic toolkit underlies terrestrial adaptation, we designed an experimental framework that directly targets these core abiotic stressors (Fig. 3a). Specifically, we exposed 17 species spanning 7 animal phyla (Nematoda, Onychophora, Arthropoda, Platyhelminthes, Mollusca, Annelida, Nemertea), to controlled stress conditions representing: desiccation and osmotic stress (water loss in terrestrial taxa; salinity shifts in aquatic taxa), oxygen variation (hypoxia and hyperoxia), UV-B radiation with recovery treatments, visible light exposure and reception of chemical cues (Fig. 1b, 3a, Supplementary Data S1, S3-S19; see Material and Methods). Following exposure, we generated 796 RNA-seq datasets across all treatments. For each species, RNA material from all experimental conditions was pooled and sequenced using IsoSeq to construct a de novo transcriptome assembly (Supplementary Data S20), which was used as a reference for mapping RNA-seq reads and gene expression quantification (Supplementary Data S3-S19). In parallel, we sequenced control samples using proteomics to validate the translation of expressed proteins.

**Figure 3.**
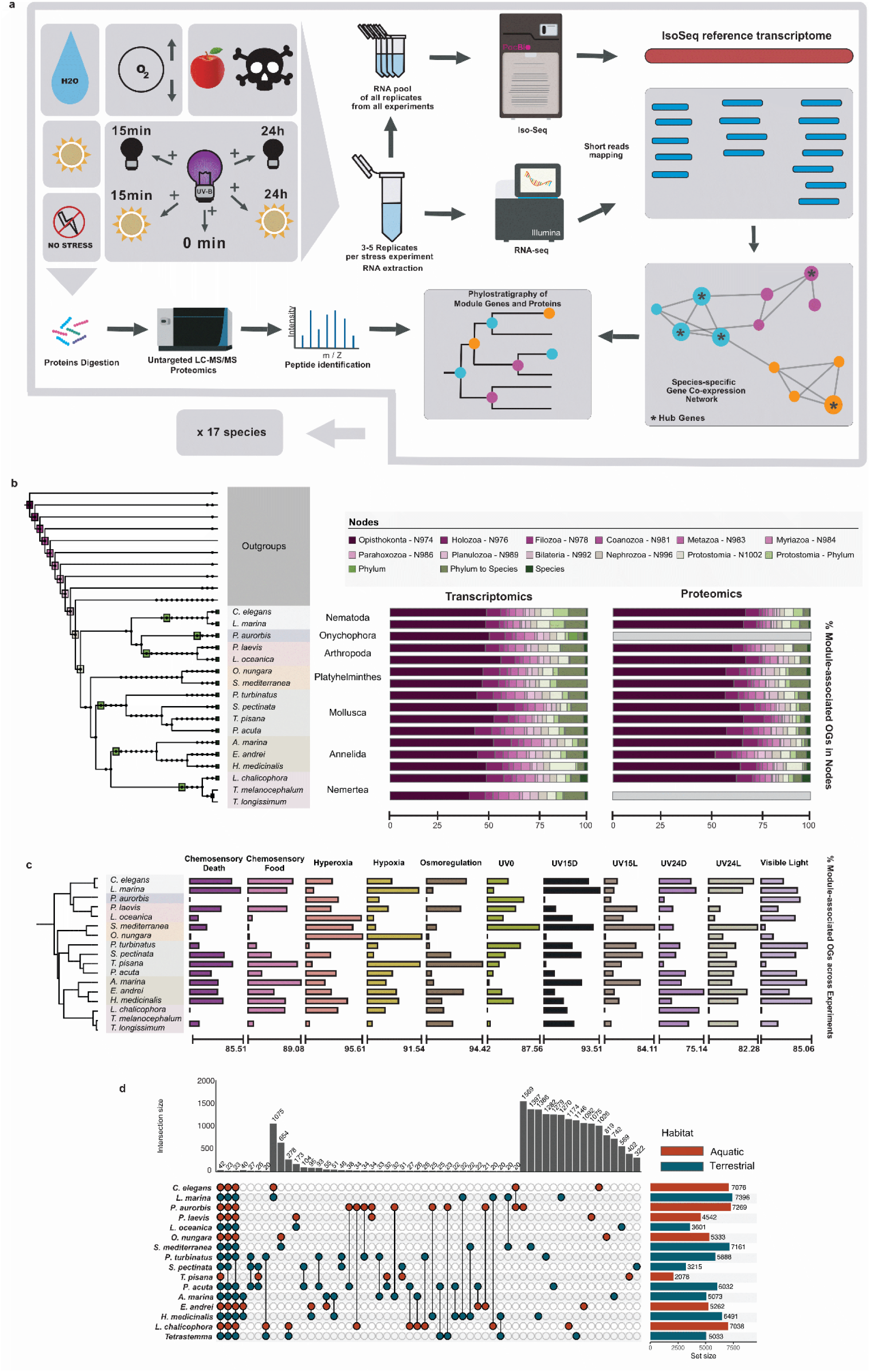
Species-specific stress-response networks and pre-metazoan evolutionary origins of stress-related genes suggest diverse genomic routes to terrestrial life in animals. **a**, Overview of abiotic stress experiments, co-expression network construction and proteomic detection of stress-related genes across 17 species from 7 animal phyla, exposed to standardised stressors mimicking terrestrial abiotic factors. **b**, Phylostratigraphic analysis showing that the majority of stress-related genes belong to old orthogroups, based on genes recovered through both transcriptomic and proteomic data. Colored squares in the tree indicate the branch leading to each node, using the same color scheme as in the accompanying barplot. Nodes from collapsed branches not shown in the tree are marked with black dots. Protostomia to Phylum and Phylum to Species segments indicate the sum of all the in-between branches. Grey bars indicate missing data for specific species. **c**, Percentage of module-associated orthogroups per species for each experiment, calculated relative to the total number of module-associated orthogroups identified in that species. **d**, UpSet plot showing overlap of module-associated orthogroups resulted from species-specific co-expression networks. Dots connect species that have a number of module-associated orthogroups in common, the number indicated in each bar from the barplot on top.

We then constructed species-specific weighted gene co-expression networks (WGCNA) based on a scale free topology (SFT) (Supplementary Fig. S94-S110, Supplementary Data S21), and identified statistically significant modules per condition (Supplementary Fig. S111-S127, Supplementary Data S22-S38). All genes belonging to these modules were retained for downstream analyses (see Data availability for gene lists) and mapped to the previously inferred orthogroups to identify their evolutionary origin (see Materials and Methods), hereafter referred to as module-associated orthogroups. In addition, we identified highly connected genes (hub genes) within each module, as these are considered to play a central role in response to stress^29–31^ (Supplementary Data S39-S55). Nonetheless, hubs are defined by network connectivity and are often enriched for essential regulatory genes and the results may be biased toward evolutionarily older components^44–47^, thus we used them as a complementary dataset to further assess the observed patterns. This framework allowed us to identify orthogroups exhibiting stress-responsive expression patterns across taxa, irrespective of their age of origin, and to test for the presence or absence of a shared genomic toolkit associated with terrestrial adaptation.

The evolutionary origin of module-associated orthogroups was determined using custom scripts based on the idea of phylostratigraphic analysis^48^ (Fig. 3b; see interactive link here: https://itol.embl.de/tree/37223118107225991770722885), enabling comparison of gene age distributions across species. To avoid redundancy, each orthogroup was counted only once per species, regardless of how many genes from that orthogroup were present within the significant modules. For unbiased cross-species comparison, we calculated the proportion of module-associated orthogroups assigned to each evolutionary branch for each species, based on the total number of module-associated orthogroups identified in that species. We repeated the analysis for genes recovered through the proteomic dataset to assess their translation into proteins. We then compared the proportion of module-associated orthogroups across all experiments and species (Fig. 3c, Supplementary Data S56) to reveal patterns in stress-response mechanisms associated with the transition to land.

Across all species, the majority of module-associated orthogroups traced back to deep evolutionary nodes, predating the emergence of terrestrial lineages and extending to early metazoan and pre-metazoan origins (Fig. 3b). Only a small fraction of module-associated orthogroups appeared to have arisen more recently, after the origin of Protostomia. While most species followed this general pattern, a few lineage-specific deviations were observed. For example, Onychophora showed an enrichment of module genes in orthogroups gained at the branch leading to the phylum as well as species-specific gains; Nematoda exhibited a moderate proportion of module-genes derived from orthogroups gained at the most recent common ancestor of Nematoidea and Panarthropoda (i.e., nematodes, nematomorphs, arthropods, tardigrades and onychophorans); Platyhelminthes showed a cluster of module-genes arising in Tricladida, consistent with prior findings^19^; and the terrestrial nemertean species, along with the terrestrial arthropod species displayed species-specific gains (see interactive link here: https://itol.embl.de/tree/37223118107225991770722885). However, even in these cases, the majority of module-genes traced back to orthogroups that predated terrestrialisation and originated in older nodes. Notably, the same pattern was consistent when we limited the analysis in the hub gene set (Supplementary Fig. S128a).

To assess whether different abiotic challenges triggered comparable module-level responses across taxa, we examined the proportion of module-associated orthogroups recovered in each species for each experimental condition (Fig. 3c, Supplementary Data S56). Although the proportion of module-associated orthogroups varied across treatments within the same species, no condition showed a consistent pattern across taxa. All stressors revealed strong responses in some species, but weak in others. This indicates that there is not a common stress-specific signature, rather than an heterogeneous, lineage-dependent employment of stress-responsive modules.

While the evolutionary origins of stress-associated genes were broadly conserved, the specific orthogroups recruited into stress-response networks were highly variable. Only 42 module-associated orthogroups out of the ca. 540,000 were shared across all 17 species (Fig. 3d), encoding proteins conserved across eukaryotes and with previously identified stress-related roles, such as RAB proteins^32^, selenoproteins^33^, ubiquitin carboxyl-terminal hydrolase 7^34^, tetraspanin^35,36^, cullin^37^, or putative voltage-dependent anion-selective channel^38^ (Supplementary Data S57). When considering the hub genes of the network, we observed virtually no overlap in the orthogroups to which hub genes belonged - not among terrestrial species, not among aquatic species, and not even within the same phylum, except for nematodes (Supplementary Fig. S128b). These findings indicate that stress responses have largely evolved through independent recruitment of distinct gene repertoires across lineages, rather than relying on a conserved genomic toolkit, though a small set of deeply conserved genes may underpin core functions across animals.

These results argue against a universal genomic toolkit for response to abiotic factors relative to life on land. The lack of orthogroup-level overlap among the module-associated genes recruited into stress-response networks suggests that different lineages independently co-opted distinct sets of genes to meet the environmental pressures associated with transition to land. Importantly, this pattern holds across both terrestrial and aquatic species and both within and between phyla, underscoring the absence of a conserved genetic solution to stress. Instead, response to abiotic stress appears to have proceeded through the lineage-specific reorganization of deeply conserved orthogroups into different genetic contexts. Overall, our findings support the idea that gene reuse diminishes with increasing evolutionary divergence^49^, even in response to similar abiotic stressors.

### Functional convergence in species-specific module genes in response to abiotic stress

In the absence of shared orthogroups among stress-response co-expression networks across species, we next asked whether convergence occurs at the level of gene function. To explore this, we applied pathway comparisons of statistically significant modules of each species and we investigated the functional similarity of hub genes identified in co-expression networks from each species exposed to abiotic stress, applying gene ontology-based comparisons, following the approach described in^18,50^ (see Methods).

First, we examined pathway-level convergence by assessing enrichment of well-defined functional categories. Despite species-specific gene repertoires, the pathways enriched in module gene sets under each stress condition were largely shared across species, indicating parallel recruitment of distinct gene sets into common functional categories. When considering all stress conditions together, a core set of conserved pathways emerged as recurrently enriched across lineages (Fig. 4a, Supplementary Fig. S129a). Pathways present in more than 80% of species included central metabolic processes (e.g., carbon and purine metabolism), genetic information processing pathways (e.g., spliceosome, ribosome, nucleocytoplasmic transport, and protein processing in the endoplasmic reticulum), as well as major signaling cascades including MAPK, PI3K–Akt, AMPK, and mTOR signaling. In addition, pathways associated with cellular stress responses (such as endocytosis, lysosome, autophagy, apoptosis, and cellular senescence) together with processes maintaining cellular architecture (focal adhesion, tight junctions, and cytoskeletal regulation) were consistently recovered across species (Fig. 4a). This recurrence of the same functional categories across phylogenetically distant species and diverse environmental conditions indicates that metazoan responses to environmental stress rely on a deeply conserved molecular toolkit. Such convergence is consistent with the concept of a universal cellular stress response, in which evolutionarily ancient pathways are repeatedly co-opted to maintain cellular homeostasis^51–53^. Nonetheless, the specific pathways activated varied depending on both the species and the type of stressor (right-side heatmaps in Fig. 4a, Supplementary Fig. S129a), suggesting that different lineages mobilize distinct components of these conserved functional modules in response to specific environmental challenges. This variation did not follow a consistent phylogenetic or ecological pattern, and is therefore likely to reflect differences in transcriptional responses (see Supplementary Figs. S130-S140 for unfiltered pathway lists per experimental condition). However, further resolution of these patterns revealed that convergence is not limited to pathway presence, but also extends to pathway prominence. The most frequently represented pathways within each experiment showed that a consistent set of pathways, including metabolic pathways, PI3K–Akt and MAPK signaling, endocytosis, focal adhesion, and components of gene expression and proteostasis, were repeatedly among the most dominant across conditions (Fig. 4b, Supplementary Fig. S129b). This indicates that stress responses are organized around a small number of core regulatory modules that are consistently prioritized across species and environmental contexts.

**Figure 4.**
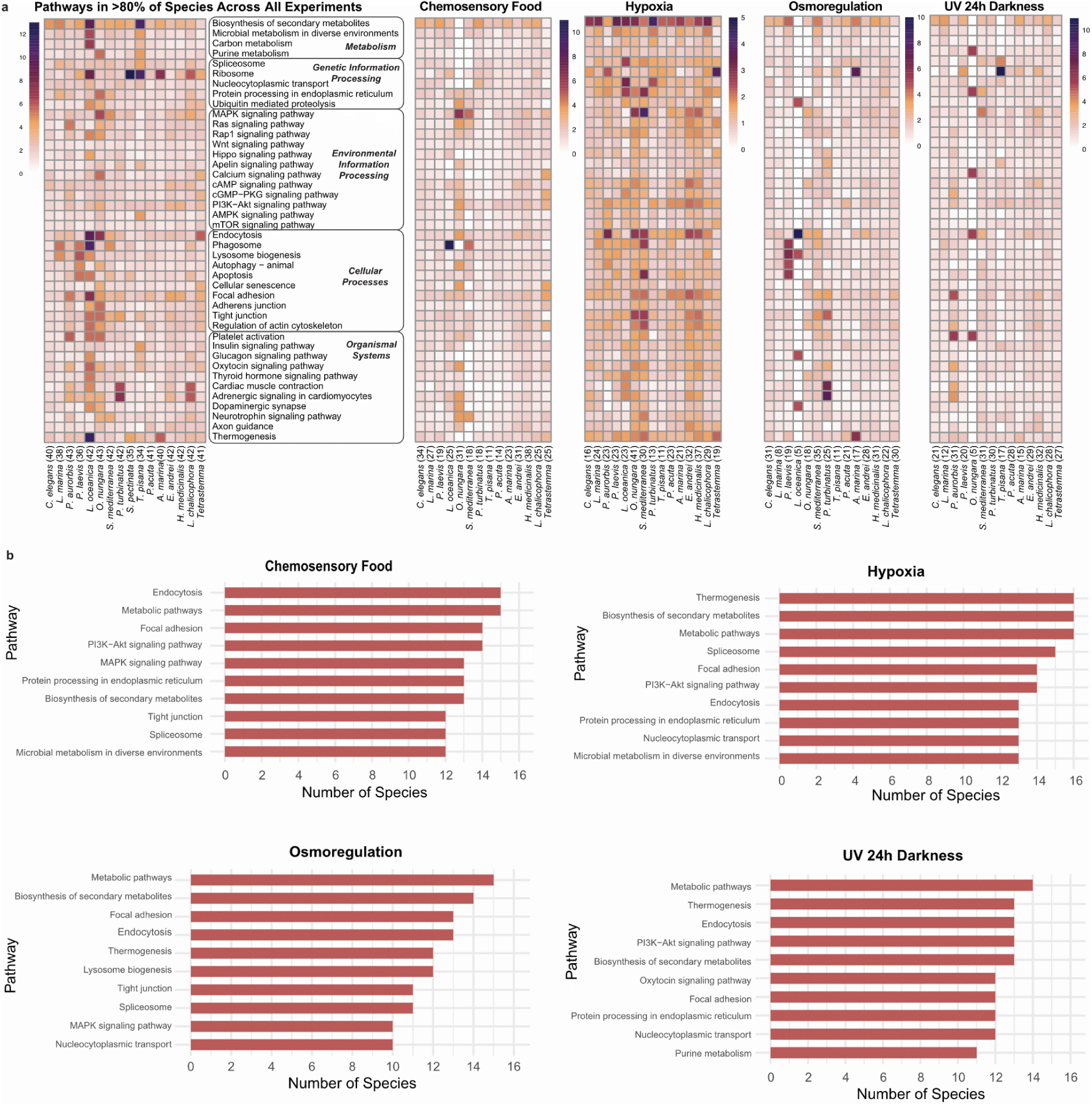
Functional convergence of species-specific KEGG pathways across animal lineages under stress conditions. **a**, Heatmaps showing the proportion of KEGG pathways in gene sets under abiotic stress conditions. Each row represents a KEGG pathway, and each column corresponds to a species. The left heatmap displays a summary heatmap including only pathways that are present in at least 80% of species, considering all experimental conditions combined. On the right, separate heatmaps are shown for each experimental condition, filtered by the same set of pathways (rows) as in the summary heatmap. Only four experiments are shown; all experiments showed the same pattern (see also Supplementary Fig. S129a). Numbers in parentheses next to species names indicate the total number of enriched pathways detected in that species for each condition. Color scales represent the percentage of genes in each pathway. **b**, Top 10 pathways per experiment. The same four experiments as in 4a are shown (see also Supplementary Fig. S129b).

To further assess the functional organization of stress-responsive modules, we examined their pathway composition across species. We found that individual modules were typically associated with multiple pathways rather than being dominated by a single functional category (Supplementary Data S58). This indicates that co-expression modules are not pathway-specific units but instead represent integrated functional assemblies encompassing multiple biological processes^54^.

To complement these results, we investigated whether core-regulatory elements (i.e., hub genes) are also showing consistent signatures of functional convergence, this time at the gene ontology (GO) level. In brief, we inferred GO annotation similarity across all hub-containing orthogroups per species, measured their functional similarity, and grouped them into functional clusters representing functional convergence units, which we represent in what we term constellation plots^18,50^ (Fig. 5a). For each experiment, we analyzed the full hub gene set of each species in an aggregated manner and we found that hub genes across different lineages shared highly similar functional annotations, though they belonged to entirely different orthogroups. When grouped by functional similarity, hub-containing orthogroups from different species clustered into one to four functional clusters including both aquatic and terrestrial animals, with most species always being included in a single main cluster (Fig. 5b, Supplementary Fig. S141). These results suggest convergent recruitment of genes with analogous roles in abiotic stress response despite divergent evolutionary origins. This pattern was further supported by orthogroup-level clustering, which revealed that the vast majority of functional clusters comprised orthogroups containing both aquatic and terrestrial species, with only a minority of clusters showing habitat-specific composition (Supplementary Data S59-S69).

**Figure 5.**
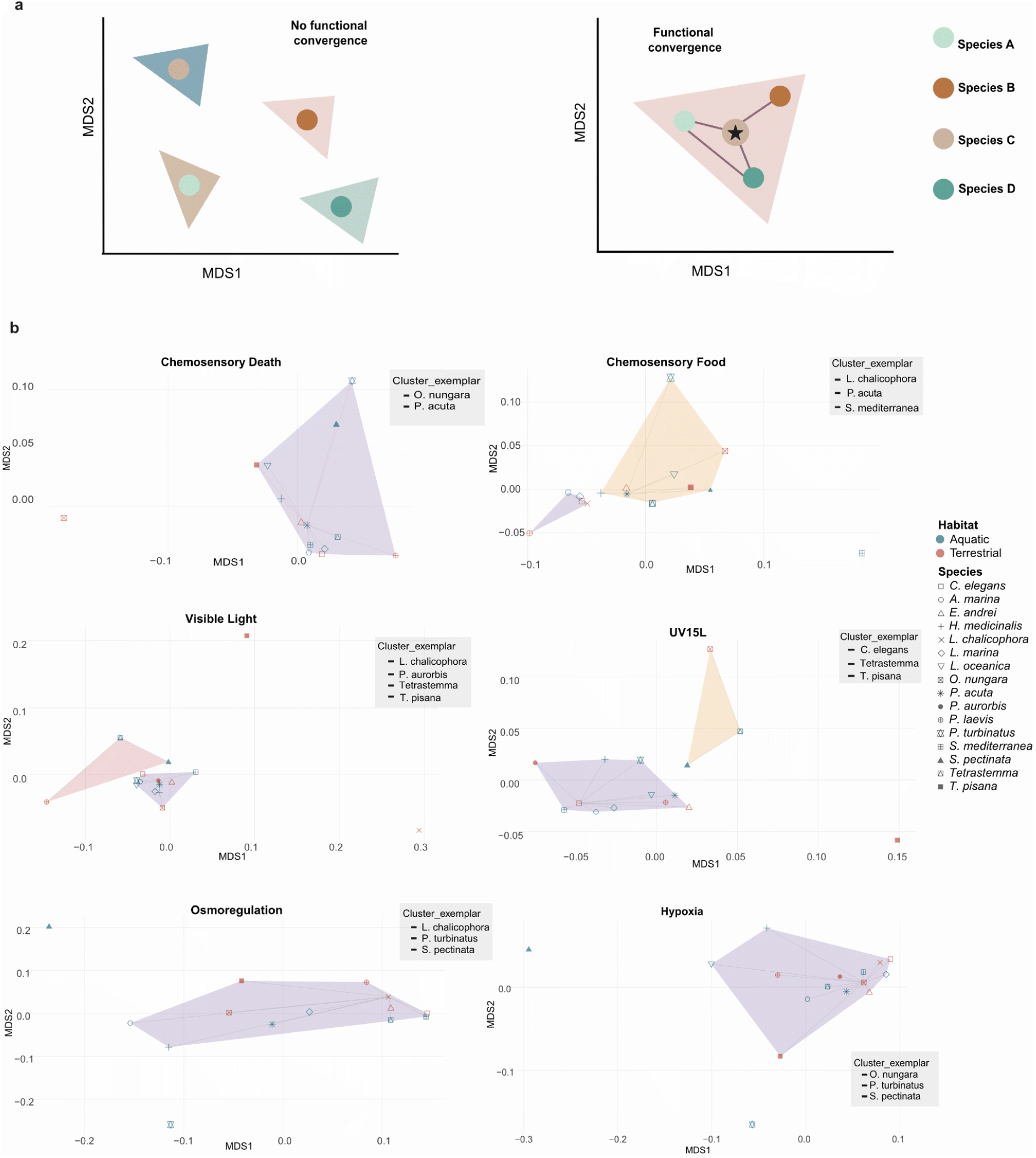
Functional convergence of species-specific, GO-annotated hub genes across animal lineages under stress conditions. **a**, Graphical illustration of constellation plots of gene functions across species. Each dot represents a species and each polygon a distinct functional cluster. On the left side each species belongs to a different cluster showing the lack of functional convergence and on the right side all species cluster together representing functional convergence. **b**, Constellation plots of functions of hub genes across 17 aquatic and terrestrial species for each experimental condition. Each polygon represents a distinct functional cluster formed by species with similar GO annotations, despite no orthogroup-level overlap across species. Cluster exemplars indicate the species that act as centroid in each cluster. The presence of hub genes from multiple species within the same functional cluster demonstrates convergence in stress-response roles across independent lineages. Only six experiments are shown for demonstration purposes, given that all experiments showed the same pattern (see also Supplementary Fig. S141). Functional convergence measured as GO-annotation similarity was inferred by calculating Wang semantic similarity between species’ GO term sets using the Best-Match Average method, followed by clustering with Affinity Propagation.

Together, these results reveal that, while the genetic content of stress-response networks differs between species, their functional roles converge across lineages. This indicates that animals transitioning to land did not reuse the same genes, but rather recruited distinct orthogroups (most of them with a pre-metazoan evolutionary origin) to produce similar functional outcomes. Such functional convergence, emerging from non-homologous gene sets, underscores the evolutionary flexibility of gene networks and suggests that response to stress during terrestrialisation was achieved through multiple genomic trajectories leading to comparable physiological outcomes.

### Machine learning identifies habitat-informative gene families across animal phyla

Although our stress-response experiments provide insight into how animals respond to abiotic challenges, they capture only part of the biological complexity involved in terrestrialisation, excluding major factors such as anatomical, embryological, biomechanical, or biochemical adaptations. To explore broader genomic signals potentially linked to the terrestrial habitat, we examined whether gene copy number variation could predict habitat. Gene copy number variation is a key mechanism of evolutionary adaptation, often reflecting functional shifts linked to ecological transitions^18,50,55^ that often underpin physiological and functional innovations. To test this, we applied machine learning, a method that has successfully identified environment-related genes in previous studies^56,57^. We trained an XGBoost machine learning classifier using our dataset of over 500,000 orthogroups inferred for our nearly 1,000 proteomes spanning all major animal phyla, providing a kingdom-wide assessment of putative habitat-associated orthogroups (Fig. 6a).

**Figure 6.**
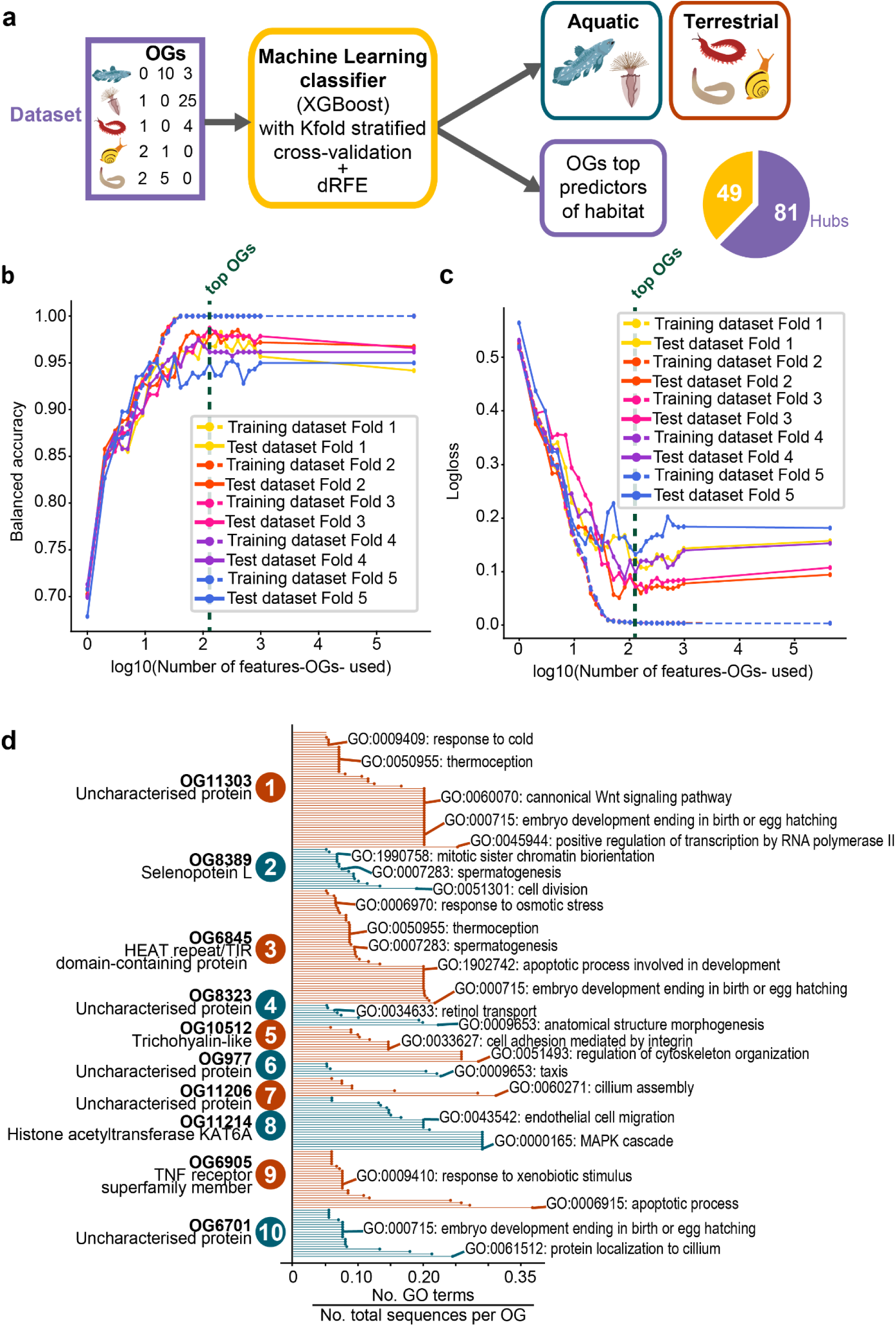
Machine learning identifies 130 orthogroups (OGs) predictors of habitat. **a**, Simplified workflow of machine learning analysis (full workflow in Supplementary Fig. S272d). The pie chart shows the number of OGs from the top 130 that contain hubs from stress-related modules in GCN. **b**, Balanced accuracy for each model trained across all iterations of the dRFE workflow for training and testing datasets of all folds. The dashed dark green line shows the iteration where we select the top OGs. **c**, Logloss values for each model trained across all iterations of the dRFE workflow for training and testing datasets of all folds. The dashed dark green line shows the iteration where we select the top OGs. **d**, Functions from the top 10 OGs as ranked by the machine learning model. OG name was obtained with BLASTp. GO terms on the right were predicted using FANTASIA. Full GO terms in Supplementary Fig. S273, Supplementary Data Table S70.

Our model identified a small subset of orthogroups (130 orthogroups, Supplementary Data S70) as the minimum number of orthogroups that accurately distinguish terrestrial from aquatic species (Fig. 6b,c). Orthogroup count distributions along the phylogeny showed that our model classified species based on complex patterns of copy number variation not easily assessed by eye (Supplementary Fig. S142-271). For example, orthogroup (OG) 8389 (Selenoprotein L, see below) is almost exclusively found in aquatic organisms (Supplementary Fig. S240), while OG492 (lysosomal acid phosphatase) shows a higher count in some terrestrial lineages than their aquatic relatives, but the opposite trend in others (Supplementary Fig. S171). This suggests that complex gene duplication and loss dynamics of genes originating before terrestrialisation events could have contributed to the transition of animals to life on land. Importantly, the discriminative power of these orthogroups does not arise from simple presence or absence or congruent expansions and contractions in independent terrestrial lineages, but from lineage-specific shifts in orthogroup size, reflecting diverse, independent, and complex evolutionary trajectories that led to convergent adaptation to terrestrial environments.

Functional annotation of these habitat-informative orthogroups revealed gene families involved in processes relevant to environmental stress, metabolism, signaling, and development (Fig. 6d, Supplementary Data S70), comprising metabolic enzymes, ion and solute transporters, and odorant receptors, which share functional roles with, or are themselves, gene families previously linked to terrestrialisation^18–22,40^, among others. Known gene families include the cytochrome P450s (CYP450s) and flavin-containing monoxygenases (FMOs), involved in the detoxification of xenobiotics, cellular metabolism, and homeostasis. Both were parallelly expanded in independent mollusc out-of-the-sea transitions^18^. CYP450s were also previously associated with critical functions during plant transition to land^58^, while FMOs are involved in the metabolism of Trimethylamine^59^, characterised by its strong fish-like odor. OG6 (serine proteases) includes vertebrate coagulation factors, one of which (factor XII) emerged before the appearance of lungfishes, during the tetrapod water-to-land transition, and has been hypothesised to confer adaptive advantages by enabling rapid clotting in soil-contaminated wounds, mitigating infection and blood loss in the terrestrial environment^60^. Other less-studied gene families identified as relevant for animal terrestrialisation were Rapunzel-like, which participates in skeletal and muscular development in fish but is absent in tetrapods^61^, and Selenoprotein L, whose function remains unknown, but was predicted in our functional analysis to be involved in chromosome segregation. The latter protein incorporates the rare amino acid selenocysteine, which depends on environmental selenium for its synthesis. It has been hypothesised that the greater bioavailability of selenium in aquatic environments, along with the increased oxidative stress associated with higher oxygen levels on land, has led aquatic species to retain larger selenoproteomes than their terrestrial counterparts^62,63^, highlighting a potential link between terrestrialisation, oxygen availability, and selenoprotein evolution. Transport of ions and solutes, such as trehalose, involved in osmoregulation, was also represented in the identified gene families. These functions are broadly consistent with the physiological demands of land life, including coping with oxidative damage, managing dehydration and oxygen variability, and processing novel chemical cues.

We further evaluated the overlap between those habitat-informative orthogroups and the hub genes previously identified in species-specific co-expression networks derived from transcriptomic experiments. Out of the 130 total orthogroups identified through machine learning, 80 contained hub genes from at least one species (Fig. 6a). This overlap suggests that gene families whose copy number variation correlates with habitat may also contribute to real-time transcriptional responses to environmental challenges. However, these orthogroups account for only a subset of the total hub gene repertoire, and their involvement as hubs is not consistent across species, even when the gene families are broadly conserved, underscoring the role of species-specific genomic rewiring in shaping stress responses.

Taken together, these results highlight a small but functionally relevant set of gene families that have undergone recurrent copy number shifts associated with habitat transitions, reflecting parallel evolutionary responses to ecological change. Moreover, their dual roles as predictive features of habitat and as transcriptional hubs suggest that these genes represent flexible genomic elements that might have been co-opted into the physiological response to stress or other challenges posed by terrestrial life.

## Discussion

The repeated transition of animal lineages from an aquatic to a terrestrial environment is one of the most impactful examples of convergent evolution. It required the independent acquisition of physiological and morphological adaptations to cope with novel environmental pressures in terrestrial habitats^28,29^. However, whether these independent adaptations might have been grounded on shared or lineage-specific genomic changes has never been addressed. Our results provide evidence that terrestrial adaptation is characterised by both convergent and parallel (different and same gene families, respectively) reshaping of terrestrial lineages’ genomic repertoires. This includes (1) independent gains and losses of genes with similar functions; (2) a lack of shared genomic responses to the same key abiotic stressors encountered during terrestrialisation, concordant with the expectation of reduced gene reuse as divergence times increase^49^, and coupled with evidence for pathway and functional convergence^64^ ; and (3) copy number variation in conserved orthogroups gained pre-terrestrialisation, primarily involved in stress responses, metabolism, and development, that might have contributed to terrestrial adaptation through complex phylogenetic patterns of duplication and loss.

Our results suggest that the evolution of terrestrial lineages relies mostly on the inherent plasticity and modularity of the pre-existing gene repertoire and molecular networks^64,68^. This contrasts with cases where adaptation relies on novel genes, such as stress response in yeasts^65^, or the repeated modification of the same genetic elements, such as the parallel loss of an enhancer causing pelvic reduction in stickleback fish^66^. Nonetheless, it is in line with the classical idea of “evolution as a tinkerer”^67^, which can be shaped by^68^ or promote the adaptation^71–73^ to the ecological pressures of terrestrial life. Previous studies have shown this trend on plant^41–43,70^, vertebrate^21^, mollusc^18^, and flatworm^19^ terrestrialisation, and on the response to desiccation in gecarcinid land crabs^69^. This, together with our results, highlights the importance of considering systems-level properties, such as co-expression network architecture, hub gene centrality, and pathway redundancy, when investigating the genomic basis of adaptation and convergent evolution^68^. Importantly, while this plasticity likely reflects intrinsic properties of ancestral gene repertoires and networks, its evolutionary outcome is expected to be context-dependent, with distinct biotic and abiotic conditions across independent terrestrialisation events shaping which components are co-opted, rewired, or lost.

Despite the breadth of our taxonomic and experimental framework, several limitations inherent to the available data and to deep-time comparative inference should be considered when interpreting our results and their generality. Firstly, although our wide taxonomic representation reduces the chance of missing information by incorporating relevant but often overlooked lineages (e.g., nemerteans), it is still biased towards lineages with a high representation in the genomic databases, limiting our confidence on the gained and lost OGs in some terrestrial lineages (e.g. our dataset only contains one terrestrial amphipod, which led us to not consider this terrestrialisation event for further analyses as we cannot distinguish between species and lineage-specific OGs). Secondly, current methods of orthology inference tend to perform less reliably when applied to highly divergent datasets or to genes with higher evolutionary rates, which can influence downstream analyses^238^. In these cases, higher sequence divergence will cause homology detection failure, creating many lineage-specific OGs that may not be real^239^ or affecting the inference of the phylostratigraphic age of origin of a gene family^240^. Moreover, these systematic errors occur on a per-gene basis, making it hard to assess the accuracy of orthology inference at a genomic scale^78^. Related to this, it should be noted that functional similarity between OGs might also reflect artifactual orthogroup oversplitting instead of distinct gene families, potentially inflating apparent functional redundancy. These caveats should be considered when interpreting the patterns obtained in this and other studies dealing with such complex large-scale datasets. Thirdly, it is not known how many terrestrialisation events occurred in animals and its inference changes depending on the species phylogeny. Perhaps the clearest example is the arachnid terrestrialisation, where the inclusion of Xiphosura within arachnids may challenge the standing idea of a single terrestrialisation event^74^, but secondary transitions to aquatic environments followed by transitions back to terrestrial habitats (e.g. in turtles^75^) can complicate the outlook even more. Further palaeontological research on invertebrates may identify additional terrestrialisation events and their ecological context, providing additional information to understand these habitat transitions. Fourthly, the separation between aquatic and terrestrial is not binary. Even though we considered terrestriality as a binary trait to make it comparable between different lineages, it could be argued that some lineages acquired more profound adaptations that allowed them to thrive in fully terrestrial habitats (e.g. arthropods that live in deserts) compared to other lineages where there is still some dependence on water (e.g. terrestrial planarians live in humid environments). A gradual classification with intermediate habitats similar to the one proposed for land crabs^76^ cannot be applied when comparing between lineages due to their profound morphological and physiological differences. Fifthly, terrestrial environments are highly heterogeneous, and the nature and intensity of abiotic stressors have likely varied substantially over evolutionary time. As such, the experimental conditions used here are best interpreted as controlled proxies for broad environmental axes (e.g. water balance, oxygen availability, UV radiation), rather than direct recreations of ancestral terrestrial settings. Furthermore, although we targeted a set of core, experimentally tractable stressors, other important factors such as temperature variation and mechanical constraints were not included and may also have contributed to terrestrial adaptation. Together, these limitations highlight that our results capture how extant taxa respond to terrestrial-like challenges, and should be interpreted as evidence of recurrent functional responses rather than direct reconstructions of ancestral molecular states.

While our present-day stress experiments and phylogenomic approach cannot recapitulate the full complexity of terrestrialisation, they provide a kingdom-wide comparative framework to understand how distantly related animal lineages achieve similar functional adaptations to terrestrial environments through the independent recruitment of distinct gene repertoires. Moreover, as computational tools and global biodiversity genomic initiatives bridge the gap between non-model and model organisms, experimental research on previously neglected lineages will become easier and will provide us with functional information to corroborate or improve computational predictions of gene function. This will benefit more in-depth studies on independent terrestrial lineages, where physiological and molecular information could be combined to get a more comprehensive view of each terrestrialisation event. All in all, our results offer a first molecular glimpse on how animals could have repeatedly overcome the challenges of novel terrestrial environments not only by inventing entirely new solutions, but mostly by tinkering pre-existing ones.

## Methods

### Species selection for kingdom-wide gene repertoire evolutionary dynamics

We selected 960 animal species covering all animal phyla, and 13 outgroup species (Supplementary Data S1). We paid special attention to their taxonomic distribution to ensure that internodes leading to terrestrial lineages were well represented. Per-gene protein sequences were obtained from MATEdb2^79^. For each species, the longest isoform per gene was parsed for orthology inference. Reference transcriptomes from species collected to carry out abiotic stress experiments were added as well (see below). Information on the source data type for each species (i.e. genome, RNA-seq transcriptome or IsoSeq transcriptome) can be found in Supplementary. Data S1. We grafted our species tree based on the information available in the literature.

### Species tree topology sources

Deep relationships between animal phyla are still the topic of debate in the scientific community. For our species selection, we decided to use the most recent and sound phylogenomic studies whenever possible (Supplementary Table S1). The use of alternative topologies to the one considered here for these unresolved relationships would not affect the general conclusions of our results, as terrestrialisation events occurred later and most orthogroups that we discussed originated before these events (regardless of the specific timing of origin).

For the most recent interrelationships, there were some cases where only a few markers or even morphological studies that include those species existed. There were a few cases where information on the exact relationship between three or four species was not available (i.e., Talitridae (Arthropoda), *Siphonaria* (Mollusca), Nerididae (Annelida), Opilioacarida (Arthropoda), Sesarmidae (Arthropoda), Hyppolytidae (Arthropoda)), forming a polytomy that was resolved using ETE v3.1.2^80^. These should not affect the conclusions, either, as our results suggest a higher importance for genes that predate these branches.

For several terrestrial lineages (e.g., Talitridae, Amphipoda), only genomic or transcriptomic data of a single terrestrial species were available. This would affect our results in the sense that we would not be able to distinguish between lineage- and species-specific orthogroups. For this reason, these cases were not considered for the gene repertoire evolution analyses.

### Extant and ancestral habitat reconstruction

Habitat information for each species was mostly obtained from the World Register of Marine Species (WoRMS) database^205^ (last accessed in January 2023). To simplify, a species was considered terrestrial if it spent at least one stage of its life cycle in a terrestrial environment. The only exceptions were the diving bell spider, *Argyroneta aquatica*, and the maritime centipede, *Strigamia maritima,* which were considered terrestrial because they conserve most adaptations to life on land.

We inferred the ancestral habitat of each branch using the R package paleotree v3.4.7^206^ using the MPR method, which provided a reconstruction more congruent with the literature than the ACCTRAN method. For ambiguous and wrongly assigned branches (Supplementary Fig. S274), we manually assigned the habitat to aquatic (Supplementary Fig. S275). Given that information on where terrestrialisation events happened is already described in the literature, and that a comprehensive testing of whether they occurred in those moments requires an even wider taxon sampling, that also incorporates fossil information (if available), and more powerful methods that cannot be applied here, the only purpose of this analysis was to map, in an automated manner, the information from the literature in the main figures. Results were not used for any additional analyses.

MPR gives ambiguous reconstructions in 3 lineages: nematodes, molluscs, and annelids. For nematodes, no paper infers the ancestral habitat state. One paper suggested more than 30 transition events (Holterman et al., 2019), but they did not distinguish between terrestrial and freshwater. The ambiguous group in question (Plectida) includes terrestrial and aquatic (marine) clades (supplementary figure 1 in that paper), making it more likely that the ancestor was aquatic. More specifically, we can say that branch 1116 from our tree is most likely aquatic, while 1117 could also be terrestrial. We used the habitats previously inferred for molluscs^180^, and annelids^197^. One species (*Drawida*) was considered as part of Crassiclitellata in their ancestral reconstruction, but not in their final selected topology. So, we considered this as a putative secondary terrestrialization event and left the ancestor as aquatic, in the absence of additional evidence that favored one of the two options.

In addition, we are also getting results inconsistent with known information from the literature. MPR considered the ancestor of all arthropods and tardigrades to be terrestrial. This was changed to aquatic.

### Gene repertoire evolutionary dynamics analyses

We used SonicParanoid2 v2.0.8 (fast mode)^207^ to reconstruct orthologous groups (orthogroups) in our 973 species. A total of ∼20M protein-coding sequences were clustered into 514,979 OGs, while ∼5M were left unclustered. OGs were used as a proxy for gene families and used to infer gene gains and losses at different nodes using ETE v3.1.2^80^. For gains, we identified the most recent common ancestor (MRCA) of all species with genes in each OG to determine their age of origin (i.e. their phylostratigraphic origin but within the phylogenetic context of the dataset). For losses, we used a modification of the custom script used in^194^ (see ’Code availability’). For each OG inferred to have originated at a given ancestral node, we assessed its presence or absence across descendant clades. A gene loss was inferred at a descendant node whenever all species descending from that node lacked genes in the OG. That is, the loss event is assigned to the MRCA of all species in which the OG was absent, provided that the OG was present in at least one species outside that clade within the parent lineage, and that the MRCA does not match the parental node. Gain-to-loss ratio was also computed at each node by dividing the number of OGs gained by the number of OGs lost.

BLASTp was run against the clustered nr database to get the identity of some OGs gained and lost. Full results are available in Figshare (see Data and Materials Availability section).

### Functional enrichment analyses

We downloaded FANTASIA v1^208^ GO functional predictions, based on similarity of embeddings from protein language models, from MATEdb2^79^ or ran new inferences for the newly IsoSeq sequenced species.

To identify the function of OGs that were gained or lost at the origin of terrestrialisation lineages (containing at least two terrestrial species), we performed a GO enrichment of the proteins contained within those OGs from selected species. For each node, we selected the smallest number of species necessary to ensure complete OG representation. For gene gains, species were selected from within the clade defined by the node. For gene losses, species were selected from the sister lineage to the clade of interest (e.g., remipedes for Hexapoda). In Myriapoda and Onychophora, where including all species was unfeasible, we ranked species by the number of OGs they contained, and selected the top 25. This subset recovered over 50% of the OGs, as shown by the cumulative increase in OGs with each added species (Supplementary Fig. S93c,d). For each terrestrialisation node, we performed separate GO enrichment analyses with topGO^209^ for gene gains and losses. In both cases, the GO enrichment analyses were first done per species using the full proteome of that species as the background. We then combined the p-values using the ordmeta method ^210^ to identify enriched GO terms for genes gained or lost at the terrestrialisation node. Functional convergence was also assessed as explained below (see semantic similarity in the ‘Functional convergence’ section).

### Stress experiments and RNA sequencing, and preprocessing

To pinpoint the gene repertoire composing the terrestrialisation genetic toolkit, we selected 17 aquatic and terrestrial species spanning 7 animal phyla (Nematoda, Onychophora, Arthropoda, Platyhelminthes, Mollusca, Annelida, Nemertea) (Fig. 1b, Supplementary Data S71). We collected and used between 19 and 68 specimens per species to carry out the abiotic stress experiments detailed below (except in nematodes, where several specimens were pooled to do the experiments due to their microscopic size) (Supplementary Data S3-S19). Animals of similar size were included in each experiment to reduce age-related effects, and individuals were randomly assigned to treatment and control groups. The species included here were selected based on contrasting habitats (aquatic and terrestrial) within each phylum, allowing us to partially control for phylogenetic background while assessing responses to stress. In some phyla (molluscs and annelids) both freshwater and marine representatives were included within each phylum in addition to terrestrial ones, to capture broader taxonomic diversity. Chordata and Tardigrada were excluded due to experimental constraints (chordates) and limited specimen availability (tardigrades). The intensity and duration of each stressor were calibrated to elicit a genetic response without causing mortality, ensuring that the conditions fell within a biologically relevant range to reveal the gene repertoire underlying abiotic stress adaptation.

***Natural light*** – Terrestrial species are regularly exposed to visible light as part of their daily environmental conditions, while many aquatic invertebrates typically experience limited or no direct light exposure. This ecological contrast suggests that light may act as a more frequent and biologically relevant stressor in terrestrial environments. To assess differential responses to visible light across aquatic and terrestrial species, we exposed individuals from all taxa to ambient natural light for 15 minutes prior to dissection and flash freezing (experiment code: VL). This treatment enabled the identification of light-induced transcriptional responses and the evaluation of lineage-specific gene expression patterns potentially associated with adaptation to terrestrial environments.

***Protection against UV radiation*** – Ultraviolet (UV-B) radiation (280–315 nm) is a potent genotoxic stressor that induces DNA damage primarily through the formation of cyclobutane pyrimidine dimers and 6–4 photoproducts^211^. These lesions cause structural distortions in the DNA helix, disrupting both transcription and replication. Two of the major repair pathways animals have evolved to mitigate such damage are: photoreactivation (light-dependent), which directly reverses DNA lesions using photolyase enzymes^212^, and nucleotide excision repair (NER, or “dark repair”), a light-independent mechanism that removes damaged nucleotides through excision and resynthesis^213^. To investigate UV-induced stress responses and the dynamics of DNA repair, specimens were exposed to UV-B radiation (302 nm) at 8.48 W/m² using a UVM-28 EL Series UV lamp (Analytik Jena US) for durations ranging from 30 seconds to 3 minutes, depending on species-specific tolerance (see Supplementary Data S72). Following exposure, animals were either flash-frozen immediately (experiment code: no recovery, UV0), or allowed to recover for 15 minutes or 24 hours under either visible light (experiment codes: UV15L, UV24L) or darkness (experiment codes: UV15D, UV24D). This design enabled us to distinguish between immediate damage responses and downstream repair mechanisms, and to experimentally separate light-dependent and light-independent pathways of DNA repair.

***Osmoregulation experiments*** – Successful transition from marine to terrestrial environments requires major physiological adaptations in water balance regulation. Among the most critical challenges for terrestrial animals is the risk of dehydration due to increased evaporative water loss in air^214^. To assess desiccation tolerance in terrestrial species, we reduced ambient relative humidity to 29–30% RH and placed individuals in petri dishes where surface moisture was removed using filter paper, a fine brush, or silica gel beads, depending on the species’ morphology. Behavioural and morphological indicators of dehydration, such as reduced activity or loss of cuticular shine, were carefully monitored, and the experiment was terminated once early signs of desiccation became apparent. In aquatic species, we examined osmoregulatory capacity by exposing marine animals to freshwater and freshwater animals to seawater for 30 seconds to 2 minutes, depending on species-specific tolerance (see Supp. Data S72 for conditions). These treatments simulate abrupt osmotic shifts and allow us to characterise gene expression responses associated with maintaining ionic and water homeostasis—key processes likely to be involved in the evolution of terrestrial life (experiment code: OR).

***Oxygen pressure experiments*** – One of the major physiological barriers to terrestrialisation is the shift from extracting oxygen in water to breathing air, which involves coping with different oxygen solubility and availability. To simulate this transition, we exposed animals to controlled oxygen extremes that reflect the lowest and highest oxygen concentrations known to support life in aquatic and terrestrial environments^215,216^. Hypoxic conditions (experiment code: Hypo) were generated using a HypoxyLab™ Hypoxia chamber (Oxford Optronix), except for Onychophora, where helium gas displacement was used as an alternative method due to logistical constraints during the field expedition. For hyperoxia (experiment code: Hyper), pure oxygen (ClearO2, UK) was sprayed into airtight glass containers until the target oxygen concentration was reached. Oxygen levels were continuously monitored using an OXY-1 SMA Single Channel Fiber Optic Oxygen Transmitter (PreSens Precision Sensing GmbH) and Planar Oxygen-Sensitive Spots affixed inside the experimental chambers. In hypoxia experiments, terrestrial animals were placed directly into airtight glass containers, which were then inserted into the HypoxyLab to achieve oxygen concentrations between 13–15% O₂. For aquatic species, the containers were pre-filled with seawater or freshwater (depending on the species), and the oxygen concentration was reduced to 8% O₂ before animals were introduced. For hyperoxia, terrestrial animals were transferred into pre-equilibrated airtight chambers after spraying with pure oxygen to achieve 40–42% O₂. Oxygen levels were maintained by additional injections as needed. In aquatic species, water was pre-oxygenated to reach 42–45% O₂ before animals were added. All exposures were conducted for 20 minutes to ensure sufficient physiological engagement without causing harm.

***Reception of chemical cues*** – The transition from aquatic to terrestrial environments necessitated profound changes in sensory systems, particularly in the detection and processing of chemical cues. While chemosensation is a universal mechanism across animals, the nature of chemical stimuli differs markedly between water and air, requiring lineage-specific adaptations in receptor types and signaling pathways. To investigate how aquatic and terrestrial invertebrates respond to ecologically relevant chemosensory cues, we designed experiments using both attractive and aversive stimuli. For attractive cues, individuals were exposed to food sources representative of their natural diet (experiment code: CF, chemosensory food) (see Supplementary Data S72 for species-specific details). To simulate warning signals, we used decomposition-associated fluids. In brief, one conspecific individual per species was euthanised and allowed to decompose in phosphate-buffered saline (PBS), freshwater, or seawater, depending on the organism’s native environment. The resulting fluid (dead conspecific) was used as an aversive chemosensory stimulus (experiment code: CD, chemosensory death). During trials, live specimens were placed individually in petri dishes (either dry or containing a small volume of water) while the stimulus (food or decomposed fluid) was introduced at the center. Behavioral responses, such as approach or avoidance, were monitored in real time. Upon a clear reaction (e.g., attraction or withdrawal), the experiment was concluded, and the sample was flash-frozen for RNA extraction.

For each treatment, we used several biological replicates per taxon and included a negative control (see Supplementary Data S3-S19 for detailed description). Specimens were flash frozen immediately after each experiment. In the case of onychophorans, samples were kept in RNAlater or TRIzol at −70°C until processed. For small animals (for instance nematodes, nemerteans and molluscs), RNA was extracted from the whole specimen. Medium-sized specimens (such as annelids or onychophorans), were dissected into anterior and posterior parts (see Supplementary Data S3-S19 for detailed description on replicates). RNA extractions after stress experiments were performed using the TRIzol® reagent (Invitrogen, USA) method following the manufacturer’s instructions and using MaXtract® High Density tubes (Qiagen) to minimize DNA contamination prior to mechanical sample homogenization either by plastic rotor pestles or by ceramic mortar, depending on the sample. The concentration of all samples was assessed by Qubit RNA BR Assay kit (Thermo Fisher Scientific). Libraries were prepared with the TruSeq Stranded mRNA library preparation kit (Illumina), and sequenced on a NovaSeq 6000 (Illumina, 2 × 150 bp) for a minimum of 6Gb coverage. Summary statistics of sequenced data are shown in Supplementary Data S3-S19.

Because most animal species do not yet have sequenced genomes, and high-quality transcriptomes have been shown to sufficiently capture gene repertoire evolutionary dynamics, particularly when completeness metrics such as BUSCO scores are high^217^, we generated full-length transcriptomes as reference datasets for co-expression network analysis. The same RNA extractions used for short-read sequencing were pooled together in each species to prepare an IsoSeq library. RNA samples were subjected to DNAse treatment using the Turbo DNA-free DNase (Invitrogen) following the manufacturer’s instructions. SMRTbell libraries were generated following the procedure ‘Preparing Iso-Seq® libraries using SMRTbell® prep kit 3.0 (PN 102-396-000 REV02 APR2022)’. To enable pooling of multiple samples on 1 SMRTcell, libraries were made using barcoded adapters. The libraries were sequenced on a Sequel-IIe using Sequel II sequencing kit 2.0 and Binding kit 3.1 with 24 hr movie-time. Summary statistics of sequenced data are shown in Supp. Data S20. For the mollusc *Siphonaria pectinata*, due to the failure to successfully prepare an IsoSeq library, we built a reference transcriptome with Trinity v2.11.0^218^ using 2 replicates per experiment condition and proceeded with the MATEdb2^79^ pipeline.

Raw Illumina RNA-seq reads for all the experiments and species mentioned above were preprocessed using Trimmomatic v0.11.9^219^ by removing adapters and ambiguous bases (MINLEN: 75, SLIDINGWINDOW: 4:15, LEADING: 10, TRAILING: 10, AVGQUAL: 30).

Both raw and trimmed reads were assessed using FastQC (https://www.bioinformatics.babraham.ac.uk/projects/fastqc) before any further analysis. For IsoSeq data, the data was demultiplexed with the IsoSeq pipeline v4.0.0 (https://github.com/PacificBiosciences/pbbioconda), and cDNA primers were removed using LIMA v2.7.1. PolyA tails and artificial concatemers were removed using isoseq refine v4.0.0, and the Full-Length Non-Concatemer (FLNC) reads were clustered into the final de-novo reference transcriptome using CD-HIT v4.8.1 (cd-hit-est, a=0.99)^220^. Each species-specific high-quality full-length reference transcriptome was evaluated using BUSCO (v5.4.7) using as reference database metazoa_odb10^221^, with all reference transcriptomes showing very high BUSCO values (Supp. Data S20), hence having sufficient quality to proceed with downstream analyses. Reference transcriptomes were translated into proteins using TransDecoder v5.5.0 (https://github.com/sghignone/TransDecoder), decontaminated using BlobTools v2 (https://blobtoolkit.genomehubs.org/blobtools2/), and the longest isoforms were obtained using a custom script for further functional analyses. The longest isoform proteomes were used as input in the orthology inference (see below).

### Gene co-expression network construction

For each species, trimmed Illumina RNA-seq reads from each experiment were quasi-mapped to the corresponding species-specific IsoSeq transcriptome using Salmon v1.10.2^222^. Genes were filtered to retain only those with ≥10 counts in at least 90% of samples across all experiments, ensuring data sparsity did not confound network construction. Expression values were transformed using the regularised log (rlog) method implemented in DESeq2 v1.44.0^223^, which stabilizes variance across expression levels and is recommended for gene co-expression analysis. Batch effects were estimated and removed using the removeBatchEffect() function from limma v3.60.6^224^, with automatic detection of hidden batch structure via surrogate variable analysis. Data integrity was assessed by identifying and excluding low-quality genes and outlier samples using the goodSamplesGenes() function from WGCNA v1.73^225^. Additional quality checks included principal component analysis and sample clustering via heatmaps (Fig. 276-292), performed using the plot_PCA() and plot_heatmaps() functions from BioNERO v1.12.0^226^, prior to network construction.

Gene co-expression networks were independently constructed for each species with Weighted Gene Co-expression Network Analysis (WGCNA) using the reads generated by stress experiments. The soft-thresholding power β was determined using the pickSoftThreshold() function, with biweight midcorrelation (bicor) and a *signed hybrid* network type to fit the scale-free topology, at the threshold of R^2^≥0.8. The mean connectivity and scale-free topology fit were visualised using softConnectivity() and scaleFreePlot() to confirm the scale-free topology of the constructed networks. Networks were constructed using the blockwiseModules() function (*signed TOMType = signed, networkType = signed hybrid, deepSplit = 4, minModuleSize =30, mergeCutHeight = 0.15*), and module color assignments were extracted for downstream analysis.

To identify significant modules and extract hub genes, we first filtered co-expression modules based on their correlation with traits of interest (stress experiments), retaining only those with significant associations (p-value < 0.05). Hub genes were then extracted by selecting genes with both high gene significance (|GS| > 0.2) and high intramodular connectivity (|KME| > 0.8) within the significant modules and used for further analysis.

### Proteomic validation of abiotic stress-induced genes

We performed untargeted LC-MS/MS proteomics to characterize proteins present in each species from those included in the abiotic stress experiments, using the control samples (see above). Samples were digested with TRIzol® reagent (Invitrogen, USA) and the protein phase was isolated, following the protocol in^227^. Samples were reduced with Tris (2-carboxyethyl) phosphine (100 mM, 60 °C, 30 min) and alkylated in the dark with iodoacetamide (20 mM, 25 °C, 30 min). The resulting protein extract was acidified with phosphoric acid. Trypsin digestion and cleanup were performed according to the S-Trap manufacturer’s instructions. Briefly, samples were loaded on S-Trap micro spin columns (ProtiFi, LLC cat # C02-micro) and washed extensively with 90% methanol containing 100 mM TEAB, pH 7.55. Trypsin was added directly to the microcolumn (1:10 w:w, 37°C, 8h, Promega cat # V5113) and then peptides were eluted by serial addition of 50 mM TEAB, 0.1% formic acid, and 0.1% formic acid in 50% acetonitrile. Protein extracts were analysed on an Orbitrap Fusion Lumos mass spectrometer coupled to an EASY-nLC 1200 system. Peptides were separated via reverse-phase chromatography over a 50-cm C18 column, using a linear acetonitrile gradient. Data were acquired in data-dependent acquisition mode with dynamic exclusion, and spectra were searched using the Mascot engine v2.6 against species-specific de novo assembled transcriptomes.

Protein identification was based on high-confidence peptide matches with a false discovery rate (FDR) of ≤1% in Protein Discoverer v2.5. Only master proteins supported by unique peptides were retained, and the homogeneity of replicates was assessed by checking the percentage of proteins present in at least 2 replicates (Supp. Data S73). Quality control was ensured via regular BSA injections and QCloud performance monitoring^228^. Further experimental details follow those described in^227^.

### Phylostratigraphic analysis of module genes

To reconstruct the evolutionary history of stress-related genes, we mapped the experimentally identified, species-specific module-associated transcripts from each condition to their corresponding proteins in the reference proteomes and assigned them to orthogroups for phylostratigraphic analysis^48^ using custom scripts (see ’Code availability’). Since species had a diverse number of module genes and these linked to varying numbers of orthogroups, we normalised the data by calculating percentages, i.e., dividing the number of genes assigned to each phylostratum (Node) by the total number of module genes per species. This approach ensured comparability across species and prevented bias due to differences in gene set sizes. The same approach was followed with module genes corroborated by proteomics. Results were visualised with iTOL v7^229^. RNA-seq data from *T. melanocephalum* and *T. longissimum* were merged at the genus level (*Tetrastemma*) because insufficient specimens were available from a single species to perform all stress experiments; as a result, different experiments were conducted on each of the two species.

The intersection of module-associated orthogroups between species and experiments was calculated and visualised with UpSet plots using the R package ComplexUpSet v1.3.3^230^ with ‘exclusive_intersection’ (i.e., distinct) mode.

Functional annotation of the 42 module-associated orthogroups shared across all species exposed to environmental stressors was performed with BLASTp against the NCBI non-redundant protein database, and KEGG pathway annotation was performed with KEGG Mapper^231^ and BlastKOALA^232^. The annotation was conducted using all the genes of *C. elegans* present in those 42 orthogroups as the representative species.

### Functional convergence

To explore potential functional convergence between genes orchestrating the transcriptomic response to the different environmental stressors in each species, we employed two strategies.

The first strategy involved identifying metabolic pathways relevant for the response to each stress in each species. To achieve this, we assigned KEGG pathways using eggNOG-mapper v2.1.12^233^ to each gene of the significant modules involved in the respective stress responses. Pathways categorised under ’Human Diseases’ and ’Drug Development’ in KEGG were excluded from further analysis. To account for differences in the number of stress-relevant genes identified per species, a pathway was considered relevant to the response if it comprised at least 1% of the total module genes for that specific species-stress context. Functional convergence was inferred by comparing the sets of relevant pathways across species and stress conditions. For visualisation, heatmaps were generated from pathway-percentage matrices, in which rows corresponded to KEGG pathways and columns to species. A summary heatmap combining all experimental conditions was constructed by retaining, for each pathway and species, the maximum observed percentage across conditions, and pathways present in more than 80% of species were retained. The same set of pathways was then used to generate condition-specific heatmaps. In all heatmaps, colour intensity represents the percentage of genes assigned to each pathway, with scales defined independently for each plot. The KEGG pathway “Metabolic pathways” was removed to avoid obscuring variation among more specific pathways. Additionally, unfiltered heatmaps were generated for each experimental condition to visualise the full pathway landscape. Module-level pathway composition was further calculated by quantifying the number of genes per module, the number of genes with KEGG annotations, the number of unique pathways represented as well as the pathway richness. Pathway richness was defined as the number of unique KEGG pathways divided by the total number of genes in the module.

The second strategy involved generating “constellation plots” using the R package constellatoR v0.1.0 (https://github.com/MetazoaPhylogenomicsLab/constellatoR) from pre-computed “Biological Process” GO terms semantic similarity matrices. We created a custom script that used pygosemsim (https://github.com/mojaie/pygosemsim; same GO release as FANTASIA: March 22^nd^, 2022) to estimate the Wang GO semantic similarity matrix between groups of GO terms using the Best-Match Average (BMA) method. Semantic similarity (*sim_BMA_*) between 2 groups (*g1* and *g2*) is calculated as follows:

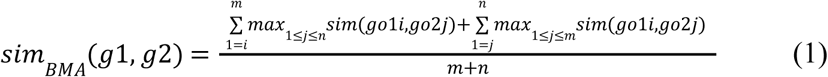

where *m* and *n* are the number of GO terms annotated in *g1* and *g2*, respectively; and *go1i* and *go2j* are the GO term in position *i* (from 1 to *m*) from the *g1* and the GO term in position *j* (from 1 to *n*) from the *g2*, respectively. sim_BMA_ semantic similarity values range from 0 to 1, with 0 meaning no similarity and 1 meaning (almost) identical.

We calculated the functional similarity between different groups: (1) enriched GO terms for gained and lost orthogroups at independent terrestrialisation events, (2) using the hub genes involved in the response to a stress condition in each species, and (3) OGs containing hub genes in the response to each stress condition.

These distance matrices were used as input for constellatoR^18,50^ to identify functional clusters, or groups exhibiting high semantic similarity, using the Affinity Propagation cluster algorithm. To visualize these results, we used scatter plots where the functional clusters are represented as polygons.

### Machine learning to identify habitat-predictive orthogroups

To identify orthogroups that may have been parallely expanded, contracted, gained, or lost in terrestrial lineages, we trained a machine learning XGBoost algorithm. We used as input the Species-OGs counts table provided by SonicParanoid2 v2.0.3a5 (outgroups removed), where we used OGs as predictive features, and the habitat of each species as the target variable. To reduce the number of features to use in the training, we pre-filtered the table to remove groups of OGs that would not be informative to predict habitat: (1) species-specific multi-copy genes, (2) OGs exclusive from fully-aquatic phyla (e.g., Porifera or Priapulida). From a total of 514,979 OGs, we ended up with 443,457 OGs after both filters.

We evaluated the performance of the model using a K-fold stratified cross-validation. Stratification (Supplementary Fig. S272a) was performed based on “phylum” (Supplementary Fig. S272b) and habitat (Supplementary Fig. S272c) to account for sampling biases in the splitting of training and test sets due to overrepresentation of specific clades or habitats in our dataset (“effect of the phylogeny”). For “phylum”, given that there were not enough species per phylum for the 5-fold cross-validation, we stratified based on the phyla that contain terrestrial and aquatic species, and for the rest, we grouped phyla into higher categories (i.e., no-Bilateria, Xenacoelomorpha, Deuterostomia, Ecdysozoa, and Lophotrochozoa).

As the number of features greatly exceeded the number of training examples in the dataset, even after pre-filtering uninformative OGs, we applied dynamic recursive feature elimination (dRFE) to iteratively reduce dimensionality based on feature importance. We used a modified implementation of Python functions from dRFEtools^234^, incorporating a strategy that combines feature importances and model accuracies across folds, as described in the original algorithm^235^.

In brief, the method begins by training the model on all features across 5 cross-validation folds. For each fold, features are ranked according to their importance in the model. These rankings are then aggregated across folds, weighted by the classification accuracy of each fold (see below), to produce an overall feature score. All features with an importance of 0 are removed in the first iteration. Then, for subsequent iterations, the bottom 30% of features (those with the lowest aggregated scores) are removed, and the process repeats with the remaining features. This continues until only one feature remains (Fig. S290d).

The ranking criterion of feature i-th is computed as follows^235^:

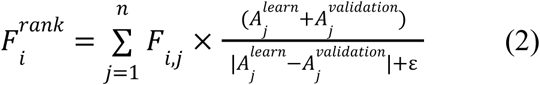

where *j*=1,.., *n* is the number of cross-validation folds, 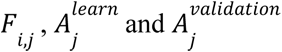 importance, the learning accuracy, and the validation accuracy of *j*-th feature, respectively. *ε* is a real number with a very small value.

From all the iterations of the dRFE, we identified the minimum number of features needed to accurately predict the environment on our species (top 130 OGs) by checking the number of features used that maximised validation accuracy and minimised log-loss in the test set (Fig. 6b,c).

To characterize those top 130 OGs (Supp. Data S70), we performed BLASTp against the nr database, and used PANGEA^236^ to do a KEGG and REACTOME pathway enrichment against *D. melanogaster, D. rerio,* and *M. musculus* references (using as input genes from those species contained in the top 130 OGs). For those OGs where there were no informative hits in any analyses, we checked the GO terms predicted by FANTASIA.

## Data Availability

The RNA-seq and IsoSeq raw data generated in this study have been deposited in the SRA NCBI database under the BioProject accession code PRJEB114751 [https://www.ebi.ac.uk/ena/browser/view/PRJEB114751]. The mass spectrometry proteomics data generated in this study are available in the ProteomeXchange Consortium via PRIDE under accession code PXD064337 [https://www.ebi.ac.uk/pride/archive/projects/PXD064337]. The data results S1 to S73 generated in this study are provided in the Supplementary Data file. The processed data for all main analyses are available at Figshare (10.6084/m9.figshare.29210009)^241^.

## Code Availability

All code and customs scripts are available at the GitHub repository associated to this manuscript https://github.com/MetazoaPhylogenomicsLab/Martinez-Redondo-Eleftheriadi_et_al_2025_G enomic_basis_animal_terrestrialization (https://doi.org/10.5281/zenodo.20700232)

## Acknowledgements

We thank Luana Da Costa Monteiro, Nicholas Stroustrup and the Planarian Lab from Universitat de Barcelona for kindly providing nematode and planarian samples to initiate the cultures used in the stress experiments; Koryu Kin and Gonzalo Berceo for letting us use their HypoxiLab for the hypoxia experiments; Salvatore Cosentino for his assistance in the installation and bug fixing of SonicParanoid2; and Leandro Aristide and Lisandra Benítez-Álvarez for sharing with us their scripts for plotting the mirror trees. We thank the three anonymous reviewers for their constructive comments, which helped us improve the manuscript. The CRG/UPF Proteomics Unit is part of the Spanish Infrastructure for Omics Technologies (ICTS OmicsTech). We also thank Centro de Supercomputación de Galicia (CESGA), particularly Pablo Rey for his assistance and guidance, and the HPC Drago from the Centro Superior de Investigaciones Científicas for access to computer resources. We declare the use of AI tools (ChatGPT, Perplexity) to polish grammar and clarity.

## Author Contributions Statement

These authors contributed equally: GIMR, KE. Author order was decided by dice rolling.

Conceptualization: GIMR, KE, RF

Specimens collection in fieldwork expeditions: FÁFÁ, KE, GIMR, JSO, IR, RGV, RF

Gene repertoire evolution analysis: GIMR

Transcriptomic and gene co-expression network analyses: KE

Stress experiments: KE, JSO, NE, RF

Wet lab experiments: JSO, NE

Machine learning analysis: GIMR, AB

Proteomics analyses: JPG, RGV, LR, CC, ES, KE

Computational analyses assistance: BC, CVC

## References

1. Selden, P. A. Terrestrialization (Precambrian-Devonian) [Internet]. eLS. (2005).

2. Vecoli, M., Clément, G. & Meyer-Berthaud, B. The Terrestrialization Process: Modelling Complex Interactions at the Biosphere-Geosphere Interface. (Geological Society of London, 2010).

3. Janvier, P. Terrestrialization: the early emergence of the concept. Geol. Soc. Spec. Publ. 339, 5–9 (2010).

4. Garwood, R. J. & Edgecombe, G. D. Early terrestrial animals, evolution, and uncertainty. Evolution (N. Y.) 4, 489–501 (2011).

5. Benton, M. J. Biodiversity on land and in the sea. Geological Journal 36, 211–230 (2001).

6. Daley, A. C., Antcliffe, J. B., Drage, H. B. & Pates, S. Early fossil record of Euarthropoda and the Cambrian Explosion. Proc Natl Acad Sci U S A 115, 5323–5331 (2018).

7. Buatois, L. A. et al. The Invasion of the Land in Deep Time: Integrating Paleozoic Records of Paleobiology, Ichnology, Sedimentology, and Geomorphology. Integr Comp Biol 62, 297–331 (2022).

8. Cloudsley-Thompson, J. L. Evolution and Adaptation of Terrestrial Arthropods. (Springer Science & Business Media, 2012).

9. de Vries, J. & Archibald, J. M. Plant evolution: landmarks on the path to terrestrial life. New Phytol 217, 1428–1434 (2018).

10. Dunlop, J. A., Scholtz, G. & Selden, P. A. Water-to-Land Transitions. Arthropod Biology and Evolution 417–439 (2013).

11. Van Straalen, N.M. Evolutionary terrestrialization scenarios for soil invertebrates. Pedobiologia 87-88, 150753 (2021).

12. Allen, K. & Briggs, D. E. G. Evolution and the Fossil Record. (1989).

13. Long, J. A. & Gordon, M. S. The Greatest Step in Vertebrate History: A Paleobiological Review of the Fish-Tetrapod Transition*. Physiological and Biochemical Zoology (2004) doi:10.1086/425183.

14. Benton, M. J. Cowen’s History of Life. (John Wiley & Sons, 2019).

15. Minter, N. J. et al. The Prelude to Continental Invasion. The Trace-Fossil Record of Major Evolutionary Events 157–204 (2016).

16. Wang, K. et al. African lungfish genome sheds light on the vertebrate water-to-land transition. Cell 184, 1362–1376.e18 (2021).

17. Meyer, A. et al. Giant lungfish genome elucidates the conquest of land by vertebrates. Nature 590, 284–289 (2021).

18. Aristide, L. & Fernández, R. Genomic insights into mollusk terrestrialization: Parallel and convergent gene family expansions as key facilitators in out-of-the-sea transitions. Genome Biol. Evol. 15, (2023).

19. Benítez-Álvarez, L. et al. Genomic exaptation and regulatory landscape shifts as key mechanisms enabling flatworm terrestrialization. Genome Biol. Evol. 18: evag063 (2026).

20. Schartl, M. et al. The genomes of all lungfish inform on genome expansion and tetrapod evolution. Nature 634, 96–103 (2024).

21. Bi, X. et al. Tracing the genetic footprints of vertebrate landing in non-teleost ray-finned fishes. Cell 184, 1377–1391.e14 (2021).

22. Martínez-Redondo, G. I. et al. Parallel duplication and loss of aquaporin-coding genes during the ‘out of the sea’ transition as potential key drivers of animal terrestrialization. Mol. Ecol. 32, 2022–2040 (2023).

23. Fleming, J. F., Pisani, D. & Arakawa, K. The Evolution of Temperature and Desiccation-Related Protein Families in Tardigrada Reveals a Complex Acquisition of Extremotolerance. Genome Biol Evol 16, (2024).

24. Little, C. The Colonisation of Land: Origins and Adaptations of Terrestrial Animals. (Cambridge University Press, 1983).

25. Willmer, P., Stone, G. & Johnston, I. Environmental Physiology of Animals. (John Wiley & Sons, 2009).

26. Cockell, C. S. Biological effects of high ultraviolet radiation on early earth--a theoretical evaluation. J Theor Biol 193, 717–729 (1998).

27. Ache, B. W. & Young, J. M. Olfaction: diverse species, conserved principles. Neuron 48, 417–430 (2005).

28. Benoit, J. B., McCluney, K. E., DeGennaro, M. J. & Dow, J. A. T. Dehydration Dynamics in Terrestrial Arthropods: From Water Sensing to Trophic Interactions. Annu Rev Entomol 68, 129–149 (2023).

29. Minelli, A., Boxshall, G. & Fusco, G. Arthropod Biology and Evolution: Molecules, Development, Morphology. (Springer Science & Business Media, 2013).

30. Fernández, R. & Gabaldón, T. Gene gain and loss across the metazoan tree of life. *Nat*. Ecol. Evol. 4, 524–533 (2020).

31. Richter, D. J., Fozouni, P., Eisen, M. B. & King, N. Gene family innovation, conservation and loss on the animal stem lineage. Elife 7, e34226 (2018).

32. Domazet-Lošo, M., Široki, T., Šimičević, K. & Domazet-Lošo, T. Macroevolutionary dynamics of gene family gain and loss along multicellular eukaryotic lineages. Nature Communications 15, 1–22 (2024).

33. Wolf, Y. I. & Koonin, E. V. Genome reduction as the dominant mode of evolution. BioEssays 35, 829–837 (2013).

34. Joyce, W. et al. A Revised Perspective on the Evolution of Troponin I and Troponin T Gene Families in Vertebrates. Genome Biol Evol 15, (2023).

35. Troyanovsky, B., Levchenko, T., Månsson, G., Matvijenko, O. & Holmgren, L. Angiomotin: an angiostatin binding protein that regulates endothelial cell migration and tube formation. J Cell Biol 152, 1247–1254 (2001).

36. Leimeister, C., Externbrink, A., Klamt, B. & Gessler, M. Hey genes: a novel subfamily of hairy- and Enhancer of split related genes specifically expressed during mouse embryogenesis. Mech Dev 85, 173–177 (1999).

37. Baba, N. & Sharma, H. M. Histochemistry of lactic dehydrogenase in heart and pectoralis muscles of rat. J Cell Biol 51, 621–635 (1971).

38. Hua, X. et al. Multi-level transcriptome sequencing identifies COL1A1 as a candidate marker in human heart failure progression. BMC Medicine 18, 2 (2020).

39. Eghbali, M., Eghbali, M., Robinson, T. F., Seifter, S. & Blumenfeld, O. O. Collagen accumulation in heart ventricles as a function of growth and aging. Cardiovasc Res 23, 723–729 (1989).

40. Brand, P. et al. The origin of the odorant receptor gene family in insects. Elife 7, e38340 (2018).

41. Dadras, A. et al. Environmental gradients reveal stress hubs pre-dating plant terrestrialization. Nat. Plants 9, 1419–1438 (2023).

42. Zhu, C. et al. WGCNA based identification of hub genes associated with cold response and development in Apis mellifera metamorphic pupae. Front. Physiol. 14, 1169301 (2023).

43. Razalli, I. I. et al. Identification and validation of hub genes associated with biotic and abiotic stresses by modular gene co-expression analysis in Oryza sativa L. Sci. Rep. 15, 8465 (2025).

44. Jeong, H., Mason, S. P., Barabási, A. L. & Oltvai, Z. N. Lethality and centrality in protein networks. Nature 411, 41–42 (2001).

45. Provero, P. Gene networks from DNA microarray data: centrality and lethality. arXiv [cond-mat.stat-mech] (2002) doi:10.48550/arXiv.cond-mat/0207345.

46. Carlson, M. R. J. et al. Gene connectivity, function, and sequence conservation: predictions from modular yeast co-expression networks. BMC Genomics 7, 40 (2006).

47. Serin, E. A. R., Nijveen, H., Hilhorst, H. W. M. & Ligterink, W. Learning from Co-expression Networks: Possibilities and Challenges. Front Plant Sci 7, 444 (2016).

48. Domazet-Loso, T., Brajković, J. & Tautz, D. A phylostratigraphy approach to uncover the genomic history of major adaptations in metazoan lineages. Trends Genet 23, 533–539 (2007).

49. Bohutínská, M. & Peichel, C. L. Divergence time shapes gene reuse during repeated adaptation. Trends Ecol Evol 39, 396–407 (2024).

50. Balart-García, P. et al. Parallel and convergent genomic changes underlie independent subterranean colonization across beetles. Nat. Commun. 14, 3842 (2023).

51. Kültz, D. Molecular and evolutionary basis of the cellular stress response. Annu Rev Physiol 67, 225–257 (2005).

52. López-Maury, L., Marguerat, S. & Bähler, J. Tuning gene expression to changing environments: from rapid responses to evolutionary adaptation. Nature Reviews Genetics 9, 583–593 (2008).

53. Hatanaka R. et al. Signaling pathways in invertebrate immune and stress response. Invertebrate Survival J. 6, 32–43 (2009).

54. Mao, L., Van Hemert, J. L., Dash, S. & Dickerson, J. A. Arabidopsis gene co-expression network and its functional modules. BMC Bioinformatics 10, 346 (2009).

55. Ishikawa, A., Yamanouchi, S., Iwasaki, W. & Kitano, J. Convergent copy number increase of genes associated with freshwater colonization in fishes. Philos. Trans. R. Soc. Lond. B Biol. Sci. 377, 20200509 (2022).

56. Davín, A. A. et al. A geological timescale for bacterial evolution and oxygen adaptation. Science 388, eadp1853 (2025).

57. Gonçalves, C. et al. Diverse signatures of convergent evolution in cactus-associated yeasts. PLOS Biology 22, e3002832 (2024).

58. Werck-Reichhart, D., Nelson, D. R. & Renault, H. Cytochromes P450 evolution in the plant terrestrialization context. Philos. Trans. R. Soc. Lond. B Biol. Sci. 379, 20230363 (2024).

59. Loo, R. L., Chan, Q., Nicholson, J. K. & Holmes, E. Balancing the equation: A natural history of trimethylamine and trimethylamine-N-oxide. J. Proteome Res. 21, 560–589 (2022).

60. Padilla, S., Prado, R. & Anitua, E. An evolutionary history of F12 gene: Emergence, loss, and vulnerability with the environment as a driver. Bioessays 45, e2300077 (2023).

61. Green, J., Taylor, J. J., Hindes, A., Johnson, S. L. & Goldsmith, M. I. A gain of function mutation causing skeletal overgrowth in the rapunzel mutant. Dev. Biol. 334, 224–234 (2009).

62. Lobanov, A. V., Hatfield, D. L. & Gladyshev, V. N. Eukaryotic selenoproteins and selenoproteomes. Biochim. Biophys. Acta 1790, 1424–1428 (2009).

63. Lobanov, A. V. et al. Evolutionary dynamics of eukaryotic selenoproteomes: large selenoproteomes may associate with aquatic life and small with terrestrial life. Genome Biol. 8, R198 (2007).

64. Hoitinga, P. K. & Birkeland, S. Pathway-level convergence: an underexplored aspect of convergent evolution. Trends Genet 41, 853–867 (2025).

65. Doughty, T. W. et al. Stress-induced expression is enriched for evolutionarily young genes in diverse budding yeasts. Nature Communications 11, 1–9 (2020).

66. Xie, K. T. et al. DNA fragility in the parallel evolution of pelvic reduction in stickleback fish. Science (2019) doi:10.1126/science.aan1425.

67. Jacob, F. Evolution and Tinkering. Science (1977) doi:10.1126/science.860134.

68. Koonin, E. V. & Wolf, Y. I. Constraints and plasticity in genome and molecular-phenome evolution. Nat Rev Genet 11, 487–498 (2010).

69. Watson-Zink, V. M., Grosberg, R. K., Lai, J. C. Y. & Bay, R. A. Transcriptomic responses of gecarcinid land crabs to acute and prolonged desiccation stress. bioRxiv 2024.07.03.601969 (2024) doi:10.1101/2024.07.03.601969.

70. Dong, Y., Krishnamoorthi, S., Tan, G. Z. H., Poh, Z. Y. & Urano, D. Co-option of plant gene regulatory network in nutrient responses during terrestrialization. Nature Plants 10, 1955–1968 (2024).

71. Helsen, J. et al. Gene Loss Predictably Drives Evolutionary Adaptation. Mol Biol Evol 37, 2989–3002 (2020).

72. Albalat, R. & Cañestro, C. Evolution by gene loss. Nat. Rev. Genet. 17, 379–391 (2016).

73. Xia, S., Chen, J., Arsala, D., Emerson, J. J. & Long, M. Functional innovation through new genes as a general evolutionary process. Nature Genetics 57, 295–309 (2025).

74. Garwood, R. & Dunlop, J. Consensus and conflict in studies of chelicerate fossils and phylogeny. Arachnologische Mitteilungen: Arachnology Letters 66:2–16 (2023).

75. Procheş, S., Polgar, G. & Marshall, D. J. K-Pg events facilitated lineage transitions between terrestrial and aquatic ecosystems. Biol Lett 10, (2014).

76. Watson-Zink, V. M. Making the grade: Physiological adaptations to terrestrial environments in decapod crabs. Arthropod Struct Dev 64, 101089 (2021).

77. Langschied, F. et al. Quest for Orthologs in the Era of Biodiversity Genomics. Genome Biol Evol 16, (2024).

78. Liebeskind, B. J., McWhite, C. D. & Marcotte, E. M. Towards Consensus Gene Ages. Genome Biol Evol 8, 1812–1823 (2016).

79. Martínez-Redondo, G. I. et al. MATEdb2, a collection of high-quality metazoan proteomes across the animal tree of life to speed up phylogenomic studies. Genome Biol. Evol. 16, evae235 (2024).

80. Huerta-Cepas, J., Serra, F. & Bork, P. ETE 3: Reconstruction, analysis, and visualization of phylogenomic data. Mol. Biol. Evol. 33, 1635–1638 (2016).

81. Ocaña-Pallarès, E. et al. Divergent genomic trajectories predate the origin of animals and fungi. Nature 609, 747–753 (2022).

82. Laumer, C. E. et al. Revisiting metazoan phylogeny with genomic sampling of all phyla. Proc. Biol. Sci. 286, 20190831 (2019).

83. Kenny, N. J. et al. Tracing animal genomic evolution with the chromosomal-level assembly of the freshwater sponge Ephydatia muelleri. Nature Communications 11, 1–11 (2020).

84. Plese, B. et al. Mitochondrial evolution in the Demospongiae (Porifera): Phylogeny, divergence time, and genome biology. Molecular Phylogenetics and Evolution 155, 107011 (2021).

85. Simion, P. et al. A Large and Consistent Phylogenomic Dataset Supports Sponges as the Sister Group to All Other Animals. Current Biology 27, 958–967 (2017).

86. DeBiasse, M. B., et al. A Cnidarian Phylogenomic Tree Fitted With Hundreds of 18S Leaves. BSSB 3, (2024).

87. Kayal, E. et al. Phylogenomics provides a robust topology of the major cnidarian lineages and insights on the origins of key organismal traits. BMC Evolutionary Biology 18, 1–18 (2018).

88. Atherton, S. & Jondelius, U. Phylogenetic assessment and systematic revision of the acoel family Isodiametridae. Zool J Linn Soc 194, 736–760 (2021).

89. Reich, A., Dunn, C., Akasaka, K. & Wessel, G. Phylogenomic analyses of Echinodermata support the sister groups of Asterozoa and Echinozoa. PLoS One 10, e0119627 (2015).

90. Cannon, J. T. et al. Phylogenomic Resolution of the Hemichordate and Echinoderm Clade. Current Biology 24, 2827–2832 (2014).

91. Zhang, Q.-L. et al. A Phylogenomic Framework and Divergence History of Cephalochordata Amphioxus. Front Physiol 9, 1833 (2018).

92. Irisarri, I. et al. Phylotranscriptomic consolidation of the jawed vertebrate timetree. Nature Ecology & Evolution 1, 1370–1378 (2017).

93. Yu, D. et al. Hagfish genome elucidates vertebrate whole-genome duplication events and their evolutionary consequences. Nature Ecology & Evolution 8, 519–535 (2024).

94. Hughes, L. C. et al. Comprehensive phylogeny of ray-finned fishes (Actinopterygii) based on transcriptomic and genomic data. Proc Natl Acad Sci U S A 115, 6249–6254 (2018).

95. Bian, C. et al. Divergence, evolution and adaptation in ray-finned fish genomes. Science China Life Sciences 62, 1003–1018 (2019).

96. Parey, E. et al. Genome structures resolve the early diversification of teleost fishes. Science 379, 572–575 (2023).

97. Hime, P. M. et al. Phylogenomics Reveals Ancient Gene Tree Discordance in the Amphibian Tree of Life. Syst Biol 70, 49–66 (2020).

98. Giribet, G. & Edgecombe, G. D. Current Understanding of Ecdysozoa and its Internal Phylogenetic Relationships. Integr Comp Biol 57, 455–466 (2017).

99. Giacomelli, M. et al. CAT-Posterior Mean Site Frequencies Improves Phylogenetic Modeling Under Maximum Likelihood and Resolves Tardigrada as the Sister of Arthropoda Plus Onychophora. Genome Biol Evol 17, evae273 (2024).

100. Herranz, M., Stiller, J., Worsaae, K. & Sørensen, M. V. Phylogenomic analyses of mud dragons (Kinorhyncha). Molecular Phylogenetics and Evolution 168, 107375 (2022).

101. Ahmed, M. et al. Phylogenomic Analysis of the Phylum Nematoda: Conflicts and Congruences With Morphology, 18S rRNA, and Mitogenomes. Front. Ecol. Evol. 9, (2022).

102. Bhat, A. H. et al. Morphological, Morphometrical and Molecular Characterization of Oscheius siddiqii Tabassum and Shahina, 2010 (Rhabditida, Rhabditidae) from India with Its Taxonomic Consequences for the Subgenus Oscheius Andrássy, 1976. Biology 10, 1239 (2021).

103. Abolafia, J. & Vecchi, M. Redescription and phylogenetic analysis of the type species of the genus Thorne, 1938 (Rhabditida, Panagrolaimidae), Thorne, 1938, including the first SEM study. J Nematol 53, (2021).

104. Bleidorn, C., Schmidt-Rhaesa, A. & Garey, J. R. Systematic relationships of Nematomorpha based on molecular and morphological data. Invertebrate Biology 121, 357–364 (2002).

105. Fleming, J. F. & Arakawa, K. Systematics of tardigrada: A reanalysis of tardigrade taxonomy with specific reference to Guil et al. (2019). Zoologica Scripta 50, 376–382 (2021).

106. Guil, N. & Giribet, G. A comprehensive molecular phylogeny of tardigrades-adding genes and taxa to a poorly resolved phylum-level phylogeny. Cladistics 28, 21–49 (2012).

107. Guil, N., Jørgensen, A. & Kristensen, R. An upgraded comprehensive multilocus phylogeny of the Tardigrada tree of life. Zoologica Scripta 48, 120–137 (2019).

108. Nichols, P. B., Nelson, D. R. & Garey, J. R. A Family Level Analysis of Tardigrade Phylogeny. Hydrobiologia 558, 53–60 (2006).

109. Stec, D., Vecchi, M., Calhim, S. & Michalczyk, Ł. New multilocus phylogeny reorganises the family Macrobiotidae (Eutardigrada) and unveils complex morphological evolution of the Macrobiotus hufelandi group. Molecular Phylogenetics and Evolution 160, 106987 (2021).

110. Stec, D., Vecchi, M., Maciejowski, W. & Michalczyk, Ł. Resolving the systematics of Richtersiidae by multilocus phylogeny and an integrative redescription of the nominal species for the genus Crenubiotus (Tardigrada). Scientific Reports 10, 1–20 (2020).

111. Baker, C. M., Buckman-Young, R. S., Costa, C. S. & Giribet, G. Phylogenomic Analysis of Velvet Worms (Onychophora) Uncovers an Evolutionary Radiation in the Neotropics. Mol Biol Evol 38, 5391–5404 (2021).

112. Costa, C. S., Giribet FLS, G. & Pinto-Da-Rocha, R. Morphological and molecular phylogeny of Epiperipatus (Onychophora: Peripatidae): a combined approach. Zool J Linn Soc 192, 763–793 (2020).

113. Giribet, G. & Edgecombe, G. D. The Phylogeny and Evolutionary History of Arthropods. Current Biology 29, R592–R602 (2019).

114. Ballesteros, J. A. et al. Comprehensive Species Sampling and Sophisticated Algorithmic Approaches Refute the Monophyly of Arachnida. Mol Biol Evol 39, msac021 (2022).

115. Xiong, Q. et al. Comparative Genomics Reveals Insights into the Divergent Evolution of Astigmatic Mites and Household Pest Adaptations. Mol Biol Evol 39, msac097 (2022).

116. Pepato, A. R., Costa, S. G. dos S., Harvey, M. S. & Klimov, P. B. One-way ticket to the blue: A large-scale, dated phylogeny revealed asymmetric land-to-water transitions in acariform mites (Acari: Acariformes). Molecular Phylogenetics and Evolution 177, 107626 (2022).

117. Ontano, A. Z. et al. Taxonomic Sampling and Rare Genomic Changes Overcome Long-Branch Attraction in the Phylogenetic Placement of Pseudoscorpions. Mol Biol Evol 38, 2446–2467 (2021).

118. Kallal, R. J. et al. Converging on the orb: denser taxon sampling elucidates spider phylogeny and new analytical methods support repeated evolution of the orb web. Cladistics 37, 298–316 (2021).

119. Wang, J., Bai, Y., Zhao, H., Mu, R. & Dong, Y. Reinvestigating the phylogeny of Myriapoda with more extensive taxon sampling and novel genetic perspective. PeerJ 9, e12691 (2021).

120. Giribet, G. & Edgecombe, G. D. Conflict between datasets and phylogeny of centipedes: an analysis based on seven genes and morphology. Proceedings of the Royal Society B: Biological Sciences (2005) doi:10.1098/rspb.2005.3365.

121. Giribet, G. & Edgecombe, G. D. Stable phylogenetic patterns in scutigeromorph centipedes (Myriapoda : Chilopoda : Scutigeromorpha): dating the diversification of an ancient lineage of terrestrial arthropods. Invert. Systematics 27, 485–501 (2013).

122. Bernot, J. P. et al. Major Revisions in Pancrustacean Phylogeny and Evidence of Sensitivity to Taxon Sampling. Mol Biol Evol 40, msad175 (2023).

123. Schwentner, M., Richter, S., Christopher Rogers, D. & Giribet, G. Tetraconatan phylogeny with special focus on Malacostraca and Branchiopoda: highlighting the strength of taxon-specific matrices in phylogenomics. Proceedings of the Royal Society B doi:10.1098/rspb.2018.1524.

124. Oakley, T. H., Wolfe, J. M., Lindgren, A. R. & Zaharoff, A. K. Phylotranscriptomics to Bring the Understudied into the Fold: Monophyletic Ostracoda, Fossil Placement, and Pancrustacean Phylogeny. Mol Biol Evol 30, 215–233 (2012).

125. Hiruta, S. F., Kobayashi, N., Katoh, T. & Kajihara, H. Molecular Phylogeny of Cypridoid Freshwater Ostracods (Crustacea: Ostracoda), Inferred from 18S and 28S rDNA Sequences. jzoo 33, 179–185 (2016).

126. Luchetti, A. et al. Comparative genomics of tadpole shrimps (Crustacea, Branchiopoda, Notostraca): Dynamic genome evolution against the backdrop of morphological stasis. Genomics 113, 4163–4172 (2021).

127. Mathers, T. C. et al. High lability of sexual system over 250 million years of evolution in morphologically conservative tadpole shrimps. BMC Evolutionary Biology 13, 1–12 (2013).

128. Copilaş-Ciocianu, D., Borko, Š. & Fišer, C. The late blooming amphipods: Global change promoted post-Jurassic ecological radiation despite Palaeozoic origin. Molecular Phylogenetics and Evolution 143, 106664 (2020).

129. Li, J.-Y., Zeng, C., Yan, G.-Y. & He, L.-S. Characterization of the mitochondrial genome of an ancient amphipod Halice sp. MT-2017 (Pardaliscidae) from 10,908 m in the Mariana Trench. Scientific Reports 9, 1–15 (2019).

130. Drumm, D. T. Phylogenetic Relationships of Tanaidacea (Eumalacostraca: Peracarida) Inferred from Three Molecular Loci. J Crustacean Biol 30, 692–698 (2010).

131. Dimitriou, A. C., Taiti, S. & Sfenthourakis, S. Genetic evidence against monophyly of Oniscidea implies a need to revise scenarios for the origin of terrestrial isopods. Scientific Reports 9, 1–10 (2019).

132. Lins, L. S. F., Ho, S. Y. W. & Lo, N. An evolutionary timescale for terrestrial isopods and a lack of molecular support for the monophyly of Oniscidea (Crustacea: Isopoda). Organisms Diversity & Evolution 17, 813–820 (2017).

133. Wägele, J.-W., Holland, B., Dreyer, H. & Hackethal, B. Searching factors causing implausible non-monophyly: ssu rDNA phylogeny of Isopoda Asellota (Crustacea: Peracarida) and faster evolution in marine than in freshwater habitats. Molecular Phylogenetics and Evolution 28, 536–551 (2003).

134. Javidkar, M., Cooper, S. J. B., King, R. A., Humphreys, W. F. & Austin, A. D. Molecular phylogenetic analyses reveal a new southern hemisphere oniscidean family (Crustacea : Isopoda) with a unique water transport system. Invert. Systematics 29, 554–577 (2015).

135. Vereshchaka, A. L., Kulagin, D. N. & Lunina, A. A. A phylogenetic study of krill (Crustacea: Euphausiacea) reveals new taxa and co-evolution of morphological characters. Cladistics 35, 150–172 (2019).

136. Wolfe, J. M. et al. A phylogenomic framework, evolutionary timeline and genomic resources for comparative studies of decapod crustaceans. Proceedings of the Royal Society B (2019) doi:10.1098/rspb.2019.0079.

137. Chak, S. T. C., Barden, P. & Baeza, J. A. The complete mitochondrial genome of the eusocial sponge-dwelling snapping shrimp Synalpheus microneptunus. Scientific Reports 10, 1–10 (2020).

138. Vereshchaka, A. L. The shrimp superfamily Sergestoidea: a global phylogeny with definition of new families and an assessment of the pathways into principal biotopes. Royal Society Open Science (2017) doi:10.1098/rsos.170221.

139. Bracken-Grissom, H. D. et al. A comprehensive and integrative reconstruction of evolutionary history for Anomura (Crustacea: Decapoda). BMC Evolutionary Biology 13, 1–29 (2013).

140. Wolfe, J. M. et al. Convergent Adaptation of True Crabs (Decapoda: Brachyura) to a Gradient of Terrestrial Environments. Syst Biol 73, 247–262 (2023).

141. Tsang, C. T. T., Schubart, C., Chu, K. H., Ng, P. K. L. & Tsang, L. M. Molecular phylogeny of Thoracotremata crabs (Decapoda, Brachyura): Toward adopting monophyletic superfamilies, invasion history into terrestrial habitats and multiple origins of symbiosis. Molecular Phylogenetics and Evolution 177, 107596 (2022).

142. Misof, B. et al. Phylogenomics resolves the timing and pattern of insect evolution. Science (2014) doi:10.1126/science.1257570.

143. Sun, X. et al. Phylomitogenomic analyses on collembolan higher taxa with enhanced taxon sampling and discussion on method selection. PLOS ONE 15, e0230827 (2020).

144. Yu, D. et al. Phylogenomics of Elongate-Bodied Springtails Reveals Independent Transitions from Aboveground to Belowground Habitats in Deep Time. Syst Biol 71, 1023–1031 (2022).

145. Yu, D. et al. Molecular phylogeny and trait evolution in an ancient terrestrial arthropod lineage: Systematic revision and implications for ecological divergence (Collembola, Tomocerinae). Molecular Phylogenetics and Evolution 154, 106995 (2021).

146. Sendra, A., Jiménez-Valverde, A., Selfa, J. & Reboleira, A. S. P. Diversity, ecology, distribution and biogeography of Diplura. Insect Conservation and Diversity 14, 415–425 (2021).

147. Carapelli, A. et al. Going Deeper into High and Low Phylogenetic Relationships of Protura. Genes 10, 292 (2019).

148. Wang, Z., Shi, Y., Qiu, Z., Che, Y. & Lo, N. Reconstructing the phylogeny of Blattodea: robust support for interfamilial relationships and major clades. Scientific Reports 7, 1–8 (2017).

149. Johnson, K. P. et al. Phylogenomics and the evolution of hemipteroid insects. Proceedings of the National Academy of Sciences 115, 12775–12780 (2018).

150. McKenna, D. D. et al. The evolution and genomic basis of beetle diversity. Proceedings of the National Academy of Sciences 116, 24729–24737 (2019).

151. Kawahara, A. Y. et al. Phylogenomics reveals the evolutionary timing and pattern of butterflies and moths. Proceedings of the National Academy of Sciences 116, 22657–22663 (2019).

152. Peters, R. S. et al. Evolutionary History of the Hymenoptera. Current Biology 27, 1013–1018 (2017).

153. Wiegmann, B. M. et al. Episodic radiations in the fly tree of life. Proceedings of the National Academy of Sciences 108, 5690–5695 (2011).

154. Khalturin, K. et al. Polyzoa is back: The effect of complete gene sets on the placement of Ectoprocta and Entoprocta. Sci Adv 8, eabo4400 (2022).

155. Lu, T.-M., Kanda, M., Satoh, N. & Furuya, H. The phylogenetic position of dicyemid mesozoans offers insights into spiralian evolution. Zoological Lett 3, 6 (2017).

156. Marlétaz, F., Peijnenburg, K. T. C. A., Goto, T., Satoh, N. & Rokhsar, D. S. A New Spiralian Phylogeny Places the Enigmatic Arrow Worms among Gnathiferans. Current Biology 29, 312–318.e3 (2019).

157. Gasmi, S. et al. Evolutionary history of Chaetognatha inferred from molecular and morphological data: a case study for body plan simplification. Frontiers in Zoology 11, 1–25 (2014).

158. Melone, G., Ricci, C., Segers, H. & Wallace, R. L. Phylogenetic relationships of phylum Rotifera with emphasis on the families of Bdelloidea. Hydrobiologia 387, 101–107 (1998).

159. Benítez-Álvarez, L. et al. Disentangling the evolutionary history of terrestrial planarians through phylogenomics. bioRxiv 2025.01.07.631701 (2025) doi:10.1101/2025.01.07.631701.

160. Laumer, C. E., Hejnol, A. & Giribet, G. Nuclear genomic signals of the ‘microturbellarian’ roots of platyhelminth evolutionary innovation. (2015) doi:10.7554/eLife.05503.

161. Orr, R. J. S. et al. Paleozoic origins of cheilostome bryozoans and their parental care inferred by a new genome-skimmed phylogeny. Sci Adv 8, eabm7452 (2022).

162. Saadi, A. J. et al. Phylogenomics reveals deep relationships and diversification within phylactolaemate bryozoans. Proceedings of the Royal Society B (2022) doi:10.1098/rspb.2022.1504.

163. Fuchs, J., Iseto, T., Hirose, M., Sundberg, P. & Obst, M. The first internal molecular phylogeny of the animal phylum Entoprocta (Kamptozoa). Mol Phylogenet Evol 56, 370–379 (2010).

164. Chen, Z. et al. A genome-based phylogeny for Mollusca is concordant with fossils and morphology. Science (2025) doi:10.1126/science.ads0215.

165. Song, H. et al. Scaphopoda is the sister taxon to Bivalvia: Evidence of ancient incomplete lineage sorting. Proceedings of the National Academy of Sciences 120, e2302361120 (2023).

166. Irisarri, I., Uribe, J. E., Eernisse, D. J. & Zardoya, R. A mitogenomic phylogeny of chitons (Mollusca: Polyplacophora). BMC Evolutionary Biology 20, 1–15 (2020).

167. Kocot, K. M., Todt, C., Mikkelsen, N. T. & Halanych, K. M. Phylogenomics of Aplacophora (Mollusca, Aculifera) and a solenogaster without a foot. Proceedings of the Royal Society B (2019) doi:10.1098/rspb.2019.0115.

168. Anderson, F. E. & Lindgren, A. R. Phylogenomic analyses recover a clade of large-bodied decapodiform cephalopods. Molecular Phylogenetics and Evolution 156, 107038 (2021).

169. Lindgren, A. R. Molecular inference of phylogenetic relationships among Decapodiformes (Mollusca: Cephalopoda) with special focus on the squid Order Oegopsida. Molecular Phylogenetics and Evolution 56, 77–90 (2010).

170. Uribe, J. E. & Zardoya, R. Revisiting the phylogeny of Cephalopoda using complete mitochondrial genomes. J Molluscan Stud 83, 133–144 (2017).

171. Uribe, J. E. et al. A Phylogenomic Backbone for Gastropod Molluscs. Syst Biol 71, 1271–1280 (2022).

172. Zhong, Z. et al. New mitogenomes in deep-water endemic Cocculinida and Neomphalida shed light on lineage-specific gene orders in major gastropod clades. Front. Ecol. Evol. 10, 973485 (2022).

173. Cunha, T. J., Reimer, J. D. & Giribet, G. Investigating Sources of Conflict in Deep Phylogenomics of Vetigastropod Snails. Syst Biol 71, 1009–1022 (2021).

174. Xu, T., Qi, L., Kong, L. & Li, Q. Mitogenomics reveals phylogenetic relationships of Patellogastropoda (Mollusca, Gastropoda) and dynamic gene rearrangements. Zoologica Scripta 51, 147–160 (2022).

175. Cunha, T. J. & Giribet, G. A congruent topology for deep gastropod relationships. Proceedings of the Royal Society B (2019) doi:10.1098/rspb.2018.2776.

176. Uribe, J. E., Colgan, D., Castro, L. R., Kano, Y. & Zardoya, R. Phylogenetic relationships among superfamilies of Neritimorpha (Mollusca: Gastropoda). Molecular Phylogenetics and Evolution 104, 21–31 (2016).

177. Yang, H. et al. Comparative Characterization of the Complete Mitochondrial Genomes of the Three Apple Snails (Gastropoda: Ampullariidae) and the Phylogenetic Analyses. International Journal of Molecular Sciences 19, 3646 (2018).

178. Colgan, D. J., Ponder, W. F., Beacham, E. & Macaranas, J. Molecular phylogenetics of Caenogastropoda (Gastropoda: Mollusca). Molecular Phylogenetics and Evolution 42, 717–737 (2007).

179. Goodheart, J. A. & Wägele, H. Phylogenomic analysis and morphological data suggest left-right swimming behavior evolved prior to the origin of the pelagic Phylliroidae (Gastropoda: Nudibranchia). Organisms Diversity & Evolution 20, 657–667 (2020).

180. Krug, P. J. et al. Phylogenomic resolution of the root of Panpulmonata, a hyperdiverse radiation of gastropods: new insight into the evolution of air breathing. Proceedings of the Royal Society B (2022) doi:10.1098/rspb.2021.1855.

181. Razkin, O. et al. Molecular phylogeny of the western Palaearctic Helicoidea (Gastropoda, Stylommatophora). Molecular Phylogenetics and Evolution 83, 99–117 (2015).

182. Doğan, Ö., Schrödl, M. & Chen, Z. The complete mitogenome of Arion vulgaris Moquin-Tandon, 1855 (Gastropoda: Stylommatophora): mitochondrial genome architecture, evolution and phylogenetic considerations within Stylommatophora. PeerJ 8, e8603 (2020).

183. González, V. L. et al. A phylogenetic backbone for Bivalvia: an RNA-seq approach. Proceedings of the Royal Society B: Biological Sciences (2015) doi:10.1098/rspb.2014.2332.

184. Smith, C. H. A High-Quality Reference Genome for a Parasitic Bivalve with Doubly Uniparental Inheritance (Bivalvia: Unionida). Genome Biol Evol 13, evab029 (2021).

185. Lemer, S., González, V. L., Bieler, R. & Giribet, G. Cementing mussels to oysters in the pteriomorphian tree: a phylogenomic approach. Proceedings of the Royal Society B: Biological Sciences (2016) doi:10.1098/rspb.2016.0857.

186. Salvi, D. & Mariottini, P. Molecular taxonomy in 2D: a novel ITS2 rRNA sequence-structure approach guides the description of the oysters’ subfamily Saccostreinae and the genus Magallana (Bivalvia: Ostreidae). Zool J Linn Soc 179, 263–276 (2016).

187. Helm, C. et al. Convergent evolution of the ladder-like ventral nerve cord in Annelida. Frontiers in Zoology 15, 1–17 (2018).

188. Li, Y. et al. Genomic adaptations to chemosymbiosis in the deep-sea seep-dwelling tubeworm Lamellibrachia luymesi. BMC Biology 17, 1–14 (2019).

189. Xu, T. et al. The Morphology, Mitogenome, Phylogenetic Position, and Symbiotic Bacteria of a New Species of Sclerolinum (Annelida: Siboglinidae) in the South China Sea. Front. Mar. Sci. 8, 793645 (2022).

190. Andrade, S. C. S. et al. Articulating ‘Archiannelids’: Phylogenomics and Annelid Relationships, with Emphasis on Meiofaunal Taxa. Mol Biol Evol 32, 2860–2875 (2015).

191. Lemer, S. et al. Re-evaluating the phylogeny of Sipuncula through transcriptomics. Molecular Phylogenetics and Evolution 83, 174–183 (2015).

192. Struck, T. H. et al. The Evolution of Annelids Reveals Two Adaptive Routes to the Interstitial Realm. Current Biology 25, 1993–1999 (2015).

193. Tilic, E., Stiller, J., Campos, E., Pleijel, F. & Rouse, G. W. Phylogenomics resolves ambiguous relationships within Aciculata (Errantia, Annelida). Molecular Phylogenetics and Evolution 166, 107339 (2022).

194. Martín-Durán, J. M. et al. Conservative route to genome compaction in a miniature annelid. *Nat*. Ecol. Evol. 5, 231–242 (2021).

195. Tilic, E., Sayyari, E., Stiller, J., Mirarab, S. & Rouse, G. W. More is needed—Thousands of loci are required to elucidate the relationships of the ‘flowers of the sea’ (Sabellida, Annelida). Molecular Phylogenetics and Evolution 151, 106892 (2020).

196. Stiller, J., Tilic, E., Rousset, V., Pleijel, F. & Rouse, G. W. Spaghetti to a Tree: A Robust Phylogeny for Terebelliformia (Annelida) Based on Transcriptomes, Molecular and Morphological Data. Biology 9, 73 (2020).

197. Erséus, C. et al. Phylogenomic analyses reveal a Palaeozoic radiation and support a freshwater origin for clitellate annelids. Zool. Scr. 49, 614–640 (2020).

198. Tong, L. et al. The genome of medicinal leech (Whitmania pigra) and comparative genomic study for exploration of bioactive ingredients. BMC Genomics 23, 1–13 (2022).

199. Andrade, S. C. S. et al. A Transcriptomic Approach to Ribbon Worm Systematics (Nemertea): Resolving the Pilidiophora Problem. Mol Biol Evol 31, 3206–3215 (2014).

200. Chernyshev, A. V., Polyakova, N. E., Norenburg, J. L. & Kajihara, H. A molecular phylogeny of Tetrastemma and its allies (Nemertea, Monostilifera). Zoologica Scripta 50, 824–836 (2021).

201. Chernyshev, A. V. & Polyakova, N. E. Distribution and Phylogenetic Position of the Antarctic Ribbon Worm Heteronemertes longifissa (Nemertea, Pilidiophora). Water 15, 809 (2023).

202. Bapst, D. W., Schreiber, H. A. & Carlson, S. J. Combined Analysis of Extant Rhynchonellida (Brachiopoda) using Morphological and Molecular Data. Syst Biol 67, 32–48 (2018).

203. Carlson, S. J. The evolution of brachiopoda. Annu. Rev. Earth Planet. Sci. 44, 409–438 (2016).

204. Hirose, M., Fukiage, R., Katoh, T. & Kajihara, H. Description and molecular phylogeny of a new species of Phoronis (Phoronida) from Japan, with a redescription of topotypes of P. ijimai Oka, 1897. Description and molecular phylogeny of a new species of Phoronis (Phoronida) from Japan, with a redescription of topotypes of P. ijimai Oka, 1897 398, 1–31 (2014).

205. WoRMS - World Register of Marine Species. https://www.marinespecies.org/aphia.php?p=popup&name=citation.

206. Bapst, D. W. paleotree: an R package for paleontological and phylogenetic analyses of evolution. Methods in Ecology and Evolution 3, 803–807 (2012).

207. Cosentino, S., Sriswasdi, S. & Iwasaki, W. SonicParanoid2: fast, accurate, and comprehensive orthology inference with machine learning and language models. Genome Biol. 25, 195 (2024).

208. Martínez-Redondo, G. I. et al. Leveraging Natural Language Processing models to decode the dark proteome across the Animal Tree of Life. Evolutionary Biology (2024).

209. TopGO. *Bioconductor* https://bioconductor.org/packages/release/bioc/html/topGO.html.

210. Yoon, S., Baik, B., Park, T. & Nam, D. Powerful p-value combination methods to detect incomplete association. Sci. Rep. 11, 6980 (2021).

211. Budden, T. & Bowden, N. A. The Role of Altered Nucleotide Excision Repair and UVB-Induced DNA Damage in Melanomagenesis. International Journal of Molecular Sciences 14, 1132–1151 (2013).

212. Weber, S. Light-driven enzymatic catalysis of DNA repair: a review of recent biophysical studies on photolyase. Biochimica et Biophysica Acta (BBA) - Bioenergetics 1707, 1–23 (2005).

213. Zachayus, A., Loup-Forest, J., Cura, V. & Poterszman, A. Nucleotide Excision Repair: Insights into Canonical and Emerging Functions of the Transcription/DNA Repair Factor TFIIH. Genes 16, 231 (2025).

214. Howard, J. M., Griffis, H.-B., Westendorf, R. & Williams, J. B. The influence of size and abiotic factors on cutaneous water loss. Advances in Physiology Education (2020) doi:10.1152/advan.00152.2019.

215. Verberk, W. C. E. P. et al. Does oxygen limit thermal tolerance in arthropods? A critical review of current evidence. Comp. Biochem. Physiol. A Mol. Integr. Physiol. 192, 64–78 (2016).

216. Mangum, C. & VAN Winkle, W. Responses of aquatic invertebrates to declining oxygen conditions. Am. Zool. 13, 529–541 (1973).

217. Cheon, S., Zhang, J. & Park, C. Is phylotranscriptomics as reliable as phylogenomics? Mol. Biol. Evol. 37, 3672–3683 (2020).

218. Grabherr, M. G. et al. Full-length transcriptome assembly from RNA-Seq data without a reference genome. Nature Biotechnology 29, 644–652 (2011).

219. Bolger, A. M., Lohse, M. & Usadel, B. Trimmomatic: a flexible trimmer for Illumina sequence data. Bioinformatics 30, 2114–2120 (2014).

220. Li, W. & Godzik, A. Cd-hit: a fast program for clustering and comparing large sets of protein or nucleotide sequences. Bioinformatics 22, 1658–1659 (2006).

221. Manni, M., Berkeley, M. R., Seppey, M., Simão, F. A. & Zdobnov, E. M. BUSCO update: Novel and streamlined workflows along with broader and deeper phylogenetic coverage for scoring of eukaryotic, prokaryotic, and viral genomes. Mol. Biol. Evol. 38, 4647–4654 (2021).

222. Patro, R., Duggal, G., Love, M. I., Irizarry, R. A. & Kingsford, C. Salmon provides fast and bias-aware quantification of transcript expression. Nat. Methods 14, 417–419 (2017).

223. Love, M. I., Huber, W. & Anders, S. Moderated estimation of fold change and dispersion for RNA-seq data with DESeq2. Genome Biol. 15, 550 (2014).

224. Ritchie, M. E. et al. limma powers differential expression analyses for RNA-sequencing and microarray studies. Nucleic Acids Res. 43, e47 (2015).

225. Langfelder, P. & Horvath, S. WGCNA: an R package for weighted correlation network analysis. BMC Bioinformatics 9, 559 (2008).

226. Almeida-Silva, F. & Venancio, T. M. BioNERO: an all-in-one R/Bioconductor package for comprehensive and easy biological network reconstruction. Funct. Integr. Genomics 22, 131–136 (2022).

227. García-Vernet, R. et al. A proteo-transcriptomic investigation of toxin evolution in planarians and their role in flatworm terrestrialization. Evolutionary Biology (2025).

228. Chiva, C. et al. QCloud: A cloud-based quality control system for mass spectrometry-based proteomics laboratories. PLoS One 13, e0189209 (2018).

229. Letunic, I. & Bork, P. Interactive Tree of Life (iTOL) v6: recent updates to the phylogenetic tree display and annotation tool. Nucleic Acids Res. 52, W78–W82 (2024).

230. Krassowski, M., Arts, M., Lagger, C. & Max. Krassowski/complex-Upset: v1.3.5. (Zenodo, 2022). doi:10.5281/ZENODO.3700590.

231. Kanehisa, M. & Sato, Y. KEGG Mapper for inferring cellular functions from protein sequences. Protein Sci 29, 28–35 (2020).

232. Kanehisa, M., Sato, Y. & Morishima, K. BlastKOALA and GhostKOALA: KEGG Tools for Functional Characterization of Genome and Metagenome Sequences. J Mol Biol 428, 726–731 (2016).

233. Cantalapiedra, C. P., Hernández-Plaza, A., Letunic, I., Bork, P. & Huerta-Cepas, J. EggNOG-mapper v2: Functional annotation, orthology assignments, and domain prediction at the metagenomic scale. Mol. Biol. Evol. 38, 5825–5829 (2021).

234. Benjamin, K. J. M., Katipalli, T. & Paquola, A. C. M. dRFEtools: dynamic recursive feature elimination for omics. Bioinformatics 39, btad513 (2023).

235. Nguyen, H.-N. & Ohn, S.-Y. DRFE: Dynamic recursive feature elimination for gene identification based on random forest. in *Lecture Notes in Computer Science* 1–10 (Springer Berlin Heidelberg, Berlin, Heidelberg, 2006).

236. Hu, Y. et al. PANGEA: a new gene set enrichment tool for Drosophila and common research organisms. Nucleic Acids Res. 51, W419–W426 (2023).

237. Holterman, M., Schratzberger, M., & Helder, J. Nematodes as evolutionary commuters between marine, freshwater and terrestrial habitats. Biological Journal of the Linnean Society, 128, 756–767 (2019).

238. Langschied, F. et al. Quest for orthologs in the era of biodiversity genomics. Genome Biol. Evol. 16, evae224 (2024).

239. Weisman, C., Murray, A. & Eddy, S. Mixing genome annotation methods in a comparative analysis inflates the apparent number of lineage-specific genes. Curr. Biol. 32, 2632–2639.e2 (2022).

240. Barrera-Redondo, J., Lotharukpong, J.S., Drost, HG. et al. Uncovering gene-family founder events during major evolutionary transitions in animals, plants and fungi using GenEra. Genome Biol 24, 54 (2023).

241. Metazoa Phylogenomics Lab (2026). Genomic basis Animal Terrestrialization. figshare. Dataset. 10.6084/m9.figshare.29210009.v1

